# Harnessing Human Immune Organoid model for Systemic and Comparative Analysis of Clinically Vaccine Adjuvants

**DOI:** 10.64898/2026.04.28.721332

**Authors:** Dandan Meng, Xuqi Li, Xiuli Rao, Xiaofen Huang, Han Ding, Qiyu Deng, Lin Li, Wentao Ma, Yuan Tao, Xiaohua Feng, Xiao Liu

## Abstract

Effective vaccine adjuvants are crucial for eliciting strong, specific, and long-lasting adaptive immune responses. However, traditional vaccine and adjuvant development is often hindered by prolonged animal testing, reliance on singular evaluation metrics, and batch variability, limiting the comparative analysis of immune responses induced by different adjuvants. To our knowledge, systematic adjuvant-focused analysis in human immune organoids remains limited, particularly across distinct antigen-conditioned immune baselines. Here, we established a matrix supported human tonsil immune organoid (M-hTIO) platform, derived from tonsil tissue for systemic adjuvant evaluation. Within 2 days, the platform replicated key germinal center (GC) features, with the potential for antigen-specific antibody production and plasmablast differentiation. Dynamic GC formation and T-B cell interactions were tracked using light-field imaging. Notably, IL-21 significantly accelerates GC-like remodeling and enhanced immune responses.

Employing bulk cell transcriptomics we systematically analyzed early immune signatures and prolonged adaptive responses induced by five clinical-grade adjuvants (Alum, MnJβ, AddaSO_3_, R848, ODN 1018) paired with influenza HA and SARS-CoV-2 RBD antigens across only adjuvant, weak and strong memory backgrounds. By integrating early and prolonged response phases, M-hTIO captured complementary dimensions of adjuvant activity, distinguishing rapid immune activation from sustained adaptive remodeling. Importantly, early activation signatures and prolonged adaptive remodeling provided complementary information, highlighting the value of temporally resolved adjuvant evaluation. AddaSO_3_ showed a consistent activity pattern across early and prolonged response phases, supporting both rapid immune activation and sustained adaptive remodeling. Moreover, our findings reveal shared/exclusive signaling pathways and gene networks. These results represent systematic and comparative analysis of clinically tested vaccine adjuvants in a human model, paving the way for high-throughput systems vaccinology research.

## Introduction

Vaccine adjuvants are critical for improving the magnitude, quality, and durability of immune responses, particularly for subunit and protein-based vaccines with limited intrinsic immunogenicity^[1–3]^.Increasing evidence indicates that vaccine outcome is strongly influenced by pre-existing immune baselines, including prior infection, vaccination history, cross-reactive exposure and pre-existing antibodies. Systems vaccinology studies have shown that some high-responder baseline states can already resemble adjuvant-induced programs^[4]^, whereas recall-immunity studies indicate that pre-existing antibodies can constrain subsequent germinal center responses^[5]^. Meanwhile, vaccine development has increasingly expanded from fixed homologous regimens toward updated boosters, heterologous immunization, and adjuvanted protein platforms^[6]^. However, most comparative studies in these settings still focus primarily on candidate efficacy, safety/reactogenicity, and terminal serological or protective outcomes, rather than systematically asking whether adjuvant choice should be tailored to distinct antigen-conditioned immune baselines^[7–9]^. Notably, a major challenge in human vaccine immunology is the lack of experimentally tractable systems that preserve the complexity of secondary lymphoid tissue^[10]^. Although blood-based analyses are practical and informative, but they incompletely reflect the evolving cellular interactions within lymphoid organs, particularly those associated with germinal center (GC) responses. What is still difficult to study directly in humans is the coupled sequence linking early activation, helper input, GC-like organization, and later differentiation output in the same tissue-relevant system. We therefore reasoned that a human immune organoid platform could be used not only to compare adjuvants, but also to test whether different formulations are preferentially effective in distinct antigen-conditioned recall states.

Human immune organoids have emerged as a promising ex vivo solution to this gap. In particular, tonsil-derived immune organoids provide access to readily available human lymphoid tissue containing diverse immune populations relevant to adaptive immunity. Wagar and colleagues^[11]^ showed that dissociated tonsil cells can reaggregate in and recapitulate key features of human adaptive immune responses. Wagoner and colleagues^[12]^ tended this platform toward systems immunology by linking organoid vaccine responses to host immune heterogeneity. In parallel, Zhong and colleagues^[13]^ further advanced engineered immune organoids to study GC biology and to model B-cell dysfunction in lymphoma-related settings. Together, these studies established that human immune organoids are biologically informative and translationally relevant. However, the key unmet need is no longer simply to show that human immune organoids can support vaccine-related testing. Rather, what remains lacking is an optimized platform better suited for quantitative and comparative immunological interrogation, particularly when the goal is to evaluate adjuvants panel across distinct antigen-conditioned immune baselines. Prior organoid studies have focused on model feasibility, donor heterogeneity, or engineered microenvironmental control than on building a structured framework for comparative adjuvant evaluation^[14]^. To our knowledge, systematic adjuvant-focused analysis in human immune organoids remains limited, especially across distinct recall-conditioned immune baselines. Additionally, organoid-compatible working concentrations remain poorly defined for many clinically relevant adjuvants. And existing organoid vaccine studies have generally proceeded directly to formulation testing without first functionally calibrating whether immune responses in the system can be directionally tuned rather than passively drifting during culture. This gap is particularly relevant for translational interpretation, where adjuvant choice may depend on the underlying recall-conditioned immune state.

Here, we addressed these gaps by establishing a matrix supported human tonsil immune organoid (M-hTIO) platform for adjuvant-oriented interrogation. We combined high-density Transwell culture with low-concentration matrix support and an improved tissue-processing workflow that increased the quality of the starting cell suspension while preserving multicellular composition. Under these conditions, the organoids reproducibly formed compact aggregates and mounted antigen-dependent GC-like responses. We defined quantitative readouts based on CXCR4^^high^ dark zone-like and CD83^^high^ light zone-like regionalization, including spatial segregation, relative marker intensity, and regional occupancy. Time-course analyses further showed that the platform supported coordinated temporal remodeling across B cells, CD4^+^T cells, and antigen-presenting compartments, indicating that the system captured evolving immune states rather than static endpoint drift. To further establish that immune responses in this platform were directionally controllable rather than passively drifting during culture, we functionally calibrated the system using biologically opposed helper-axis perturbations, IL-21 supplementation and ICOS-L blockade. Live imaging further enabled direct visualization of T/B cells lymphocyte redistribution during GC-like organization in the human organoid context.

We then applied this optimized and functionally validated platform to a panel of clinical adjuvants, including aluminum (Al) adjuvant, AddaSO_3_, ODN1018, R848, and MnJβ. To evaluate formulation behavior across distinct immune baselines, we established antigen-conditioned organoid cohorts using two model antigens, influenza HA and SARS-CoV-2 RBD. Using this framework, we organized the platform into three cohorts: a weak-pre-existing memory organoid cohort (AO) used for adjuvant-prime stimulation, a weak pre-existing memory cohort (WPO) stimulated with influenza HA, and a strong pre-existing memory cohort (SPO) stimulated with SARS-CoV-2 RBD. After defining organoid-compatible working concentrations, we evaluated these formulations with two model antigens, SARS-CoV-2 RBD and influenza HA, across distinct antigenic recall settings and across early and prolonged response phases. This design enabled us to ask not only which formulations rapidly trigger immune activation, but also which responses are more likely to be maintained across time and across biologically distinct recall backgrounds. By integrating phenotypic readouts with bulk transcriptomic profiling, our study establishes optimized M-hTIO as a human tissue-relevant platform for baseline-aware comparative adjuvant evaluation, prioritizing candidate formulations in a human tissue-relevant setting.

## Establishment of a human tonsil immune organoid platform with reproducible GC-like responses

Previous studies have established immune organoids generated from dissociated tonsil immune cells on permeable supports^[11]^, and epithelial organoids generated from matrix-embedded tonsil epithelial cells^[15]^. Building on these approaches, we established a human tonsil immune organoid platform by combining high-density culture on transwells with low-concentration matrix support (**Fig. 1, see Supplementary Table 1 for tissue donor characteristics)**. Hereafter, we refer to this system as M-hTIO.

**Figure 1.**
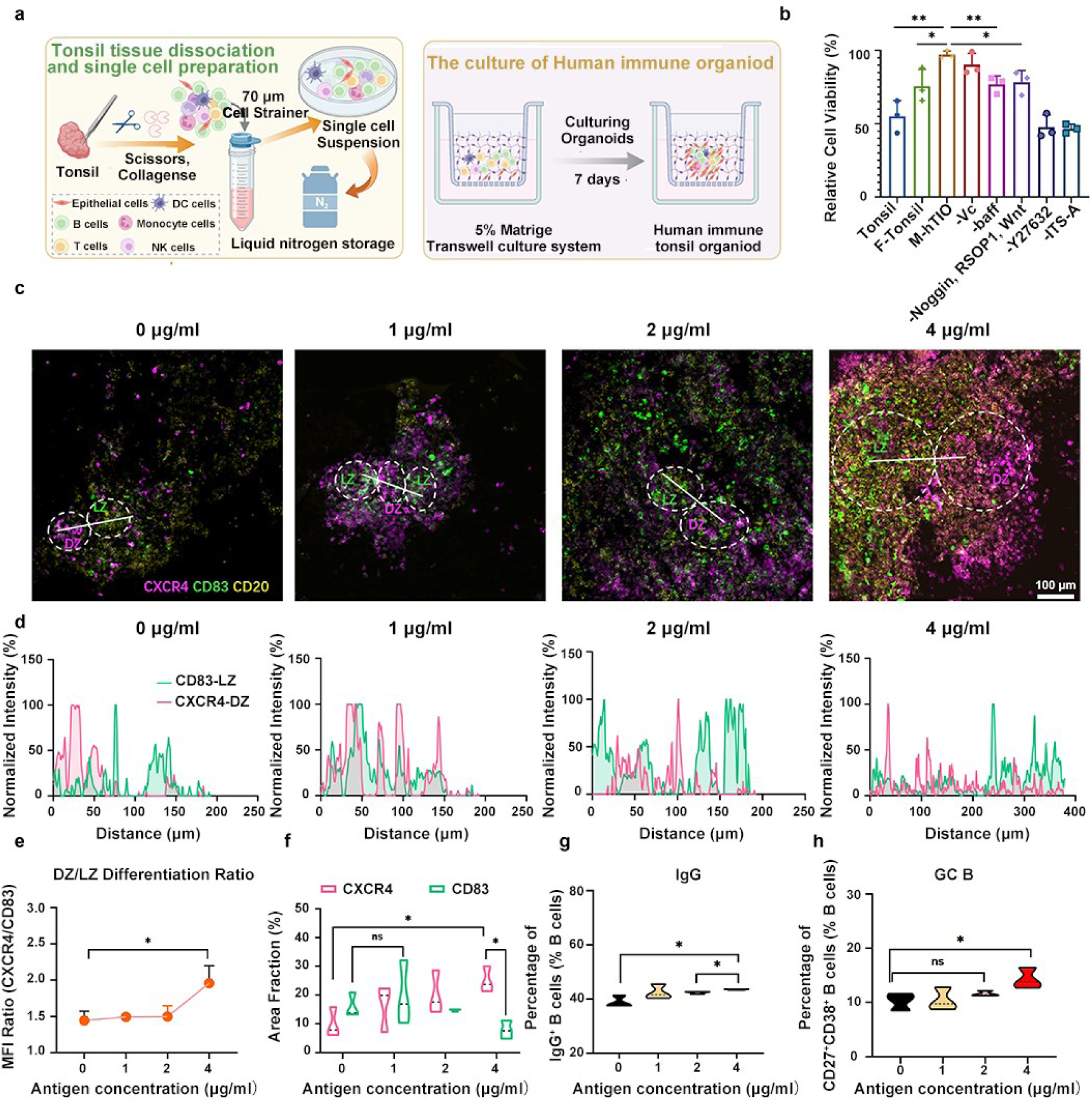
Establishment of a human tonsil immune organoid platform with reproducible GC-like responses. **a**, Schematic overview of the M-hTIO organoid workflow. Dissociated tonsil cells were cultured at high density on transwells with low-concentration matrix support under defined medium conditions. A staged tissue-processing workflow was used before organoid culture. **b,** Comparison of culture media for transwell tonsil organoid maintenance. Relative viability after 7 days of culture is shown for the indicated media conditions. **c,** Representative confocal images of organoids stimulated for 7 days with 0, 1, 2, or 4 μg/mL SARS-CoV-2 NTD. CXCR4^high regions were defined as DZ-like regions and CD83^high regions as LZ-like regions. **d,** Representative fluorescence line-scan analysis showing spatial distribution of CXCR4 and CD83 signals within GC-like regions. **e,** Quantification of the CXCR4-DZ/CD83-LZ intensity ratio across antigen doses, measured in three representative GC-like regions per condition. **f,** Quantification of the relative area occupied by CXCR4-high DZ-like and CD83-high LZ-like regions within the GC-like ROI, measured in the three representative GC-like regions per condition. Percent area values for DZ-like and LZ-like regions were calculated independently and therefore were not constrained to sum to 100%. g–h, Flow cytometric analysis of GC-associated B-cell differentiation states after 7 days of antigen stimulation, including IgG^+^ B cells and GC B cells. Together, these results establish a M-hTIO platform that supports reproducible reaggregation and quantifiable antigen-dependent GC-like responses. Data are shown as mean ± SEM with individual donors overlaid. For comparisons between two groups, paired or unpaired two-tailed Student’s t-tests were used as appropriate. For experiments with more than two groups, statistical analysis was performed using two-way repeated-measures ANOVA followed by Tukey’s multiple comparisons test. ns, not significant; *P < 0.05, **P < 0.01, ***P < 0.001, ****P < 0.0001.

Before organoid culture, we established a staged tissue-processing workflow that removed necrotic tissue and incorporated decontamination and antimicrobial pre-treatment. A short initial enzymatic digestion followed by gentle mechanical dissociation released most lymphocytes with reduced procedure-related damage, whereas continued digestion of the remaining tissue recovered epithelial, stromal-associated, and residual lymphoid cells. This workflow increased the viability of the resulting single-cell suspension from 74.6% to 91.9% **(Supplementary Fig. S1a)**. Flow cytometric analysis further showed that the optimized procedure preserved a lymphocyte-dominant population while retaining a measurable epithelial fraction **(Supplementary Fig. S1b)**, thereby providing a suitable cellular foundation for subsequent tonsil immune organoid generation.

We next defined culture conditions for organoid maintenance. Two previously reported tonsil organoid media, Tonsile^[11]^ and F-Tonsile^[15]^, were compared, and their compositions were further refined through a factor-withdrawal strategy **(Fig. 1b)**. The tested formulations included supportive factors such as Noggin, vitamin C, BAFF, Wnt, RSPO1, Y27632, and ITS-A. Among all tested conditions, the Final medium maintained the highest relative viability after 7 days of transwell culture, showing higher viability than multiple factor-depletion conditions **(Fig. 1b)**. Morphologically, both Final and F-Tonsile supported the formation of more organoid-like aggregates than in Tonsile medium **(Fig. S1c)**. Given primary organoid cultures are often sensitive to serum source, we further compared several commonly used and commercially available serum brands, including LONSERA, VisionEstern, Biochannel, ExCell, and Gibco. Gibco showed better cell viability overall and was therefore selected for subsequent experiments, whereas Biochannel and ExCell performed comparably under the tested conditions (**Supplementary Fig. S1d**). To provide additional structural support and a supportive matrix environment during transwell culture, we supplemented the system with low-concentration Matrigel. Pilot experiments showed that 3%–5% Matrigel better preserved aggregate structure than higher matrix concentrations, and 5% Matrigel was therefore selected for subsequent experiments (**Supplementary Fig. S1e)**. We used SARS-CoV-2 NTD as a model antigen to evaluate whether the platform could mount a quantifiable antigen-dependent GC-like response. Under the final culture conditions, dissociated tonsil cells progressively reaggregated over 7 days and formed compact organoid-like clusters, indicating good reproducibility across donors (**Supplementary Fig. S2**). Gross aggregate morphology was similar between unstimulated and antigen-stimulated cultures under bright-field imaging. We next asked whether antigen stimulation induced a quantifiable GC-like response within these reaggregated cultures. Organoids were stimulated for 7 days with 0, 1, 2, or 4 μg/mL NTD antigen and then examined by confocal imaging. GC-like organization was detectable across all tested doses; however, increasing antigen concentration resulted in progressively clearer spatial segregation between DZ like (CXCR4^^high^) and LZ like (CD83^^high^) regions (**Fig. 1c**). In this study, CXCR4^high regions were defined as DZ-like regions, whereas CD83^high regions were defined as LZ-like regions, providing a framework for quantifying GC-like zonation within the organoid model. To quantify this organization, we assessed three complementary features of GC-like maturation, including spatial segregation, functional polarization, and regional restructuring (**Fig. 1d–f**). First, a representative line-scan across GC-like region showed that CXCR4 and CD83 largely overlapped in the unstimulated group, but were separated by ∼50 μm after 4 μg/mL antigen stimulation, indicating clearer DZ-like/LZ-like organization (**Fig. 1d**), indicating a clearer DZ-like/LZ-like spatial segregation. Second, the CXCR4-DZ/CD83-LZ intensity ratio was used to evaluate the relative dominance of DZ-like versus LZ-like programs. Under low-dose stimulation (0–2 μg/mL), the CXCR4-DZ/CD83-LZ intensity ratio remained relatively stable, whereas 4 μg/mL antigen increased the ratio to 1.958 ± 0.242 (**Fig. 1e**), indicating a stronger DZ-like bias at the highest antigen dose. Third, we quantified the relative area occupied by CXCR4-high DZ-like and CD83-high LZ-like regions within the GC-like ROI (**Fig. 1f**). Under control conditions, the CXCR4-high and CD83-high regions occupied 9.930 ± 5.185% and 16.011 ± 4.281% of the GC-like ROI, respectively. In contrast, 4 μg/mL antigen increased the relative area of the CXCR4+ DZ-like region to 25.022 ± 4.497% while reducing the CD83+ LZ-like region to 7.873 ± 3.164% (**Fig. 1f**). Together, these results indicate that high-dose antigen stimulation promoted not only stronger GC marker expression, but also a more polarized and spatially compartmentalized GC-like architecture.

Flow cytometric analysis further supported the imaging results. Although total T-cell and B-cell frequencies were not substantially altered across antigen doses (**Supplementary Fig. S3a, Fig. S3f**), increasing antigen concentration was associated with higher frequencies of IgG^+^ B cells, GC B cells, and pre-GC B cells (**Fig. 1g–h, Fig. S3b, Fig. S3h**). By contrast, naive B cells and memory B cells decreased (**Fig. S3c-d)**, whereas plasmablasts showed an upward trend without reaching statistical significance (**Supplementary Fig. S3e, Fig. S3g**). Together, these findings suggest that antigen dose primarily reshaped the distribution of B-cell differentiation states within the M-hTIOs, rather than altering gross lymphocyte composition. Based on the combined confocal and flow cytometric data, 4 μg/mL antigen was selected as the standard working concentration for subsequent experiments.

## Temporal dynamics of innate and adaptive immune responses in tonsil organoids

We next asked whether tonsil immune organoids undergo time-dependent immune remodeling during culture, and how antigen stimulation reshapes selected components of this process. To address this, organoids were cultured in the presence or absence of NTD antigen and analyzed at days 0, 3, and 7, aiming to characterize time-dependent remodeling of lymphoid and antigen-presenting cell compartments within the system. (**Fig. 2; Supplementary Fig. S4-S5**). Representative CD27–CD38 quadrant plots from one donor are shown in **Fig. 2a**, illustrating progressive redistribution of B-cell states between days 0, 3, and 7 in both unstimulated and antigen-stimulated organoids. Group-level quantification showed that naive B cells declined over time in both conditions, with a more evident overall reduction in the antigen-stimulated group by day 7 (**Fig. 2b**). Donor-level trajectories were broadly consistent with this pattern, with most samples showing a downward trend over time (**Supplementary Fig. S4a**). In contrast, GC B cells increased during culture and showed a clearer late rise under antigen stimulation, particularly between days 3 and 7 (**Fig. 2c, Supplementary Fig. S4b**). Plasmablasts also increased over time in both groups and became more abundant at later time points (**Fig. 2d, Supplementary Fig. S4c**), suggesting progression of a fraction of B cells toward antibody-secreting differentiation. Pre-GC B cells declined over time in both groups, consistent with a transitional state during ongoing B-cell remodeling (**Fig. 2e**). At the donor level, however, trajectories were more heterogeneous, with a subset of samples showing a transient rise at day 3 before decreasing by day 7 (**Supplementary Fig. S4d**). Memory B cells increased during culture, but no clear separation between unstimulated and antigen-stimulated organoids was observed at day 7, suggesting that this change mainly reflects time-dependent remodeling within the culture system rather than a strong stimulation-specific endpoint effect (**Supplementary Fig. S4e-f**). Together, these data indicate that tonsil immune organoids undergo intrinsic temporal remodeling of B-cell states during culture, while antigen stimulation most clearly reinforces the late emergence of GC-associated populations.

**Figure 2.**
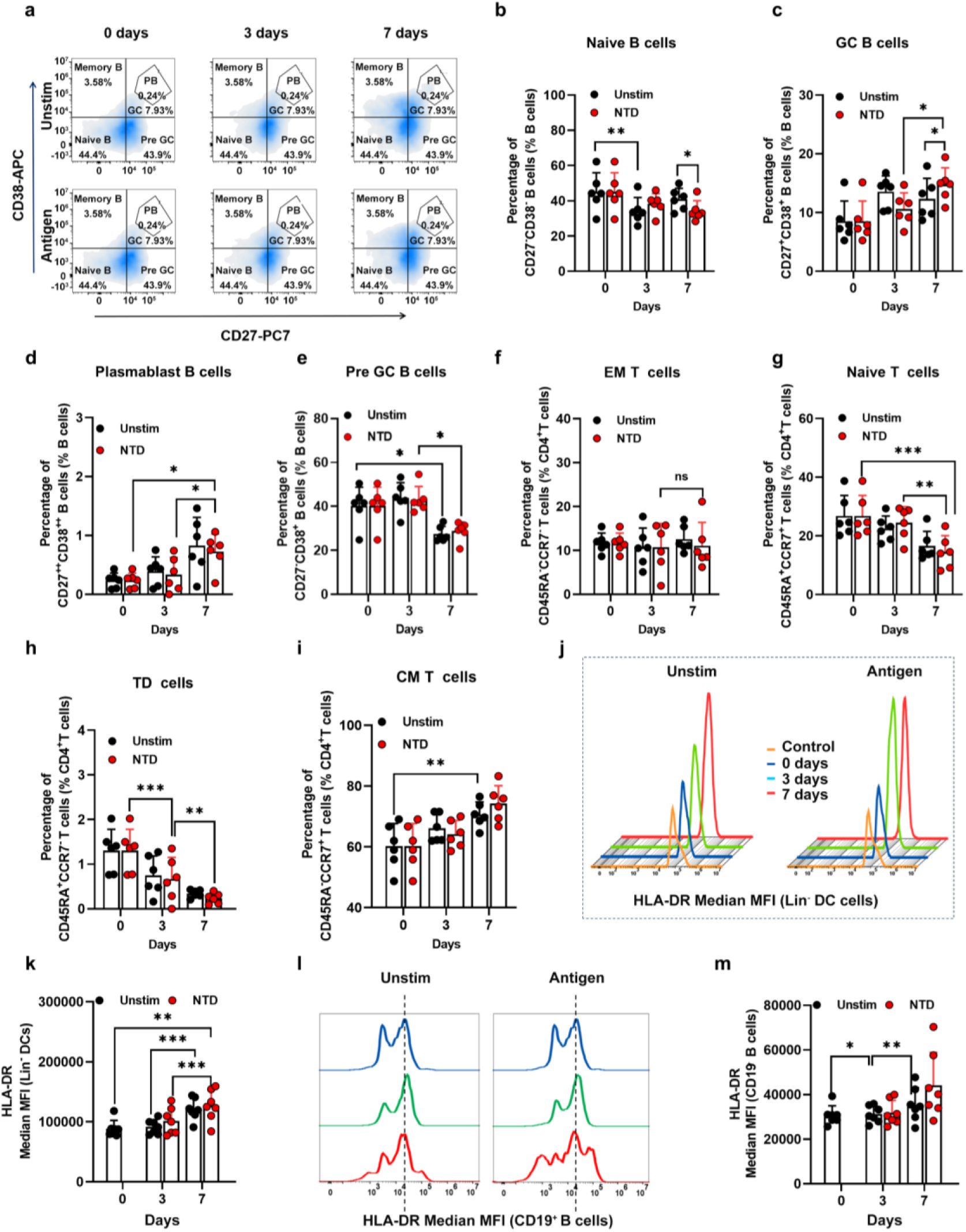
Temporal remodeling of B cell, T cell, and antigen-presenting compartments in tonsil immune organoids under unstimulated and antigen-stimulated conditions. **a**, Representative CD27–CD38 quadrant plots from one donor at days 0, 3, and 7 under unstimulated and NTD-stimulated conditions, showing temporal redistribution of B-cell states during culture. **b–e,** Quantification of major B-cell subsets over time, including naive B cells (**b**), GC B cells (**c**), plasmablast B cells (**d**), and pre-GC B cells (**e**). **f–i,** Quantification of CD4⁺T cell subsets over time, including EM T cells (**f**), TD T cells (**g**), CM T cells (**h**), and naive T cells (**i**). **j,** Representative overlaid histograms showing HLA-DR expression in Lin⁻ DC-like cells at days 0, 3, and 7 under unstimulated and antigen-stimulated conditions. Lin⁻ DC-like cells were defined as CD45⁺CD19⁻CD3⁻CD56⁻CD14⁻HLA-DR⁺ cells. **k**, Quantification of HLA-DR median fluorescence intensity (MFI) in Lin⁻ DC-like cells over time. **l,** Representative overlaid histograms showing HLA-DR expression in CD19⁺ B cells at days 0, 3, and 7 under unstimulated and antigen-stimulated conditions. **m,** Quantification of HLA-DR median fluorescence intensity (MFI) in CD19⁺ B cells over time. (n = 6), data are shown as mean ± SEM with individual donors overlaid. Statistical analyses were performed using two-way repeated-measures ANOVA followed by Tukey’s multiple comparisons test. ns, not significant; *P < 0.05, **P < 0.01, ***P < 0.001, ****P < 0.0001.

Because GC-associated B-cell remodeling is expected to occur in concert with helper T-cell support, we next examined temporal changes in the T-cell compartment (**Supplementary Fig. S5)**. Across the CD4⁺T compartment, temporal remodeling was evident during culture, although the magnitude and direction of change differed across subsets. Effector memory (EM) CD4⁺T cells showed only modest fluctuation over time and did not exhibit a clear late accumulation or consistent separation between unstimulated and antigen-stimulated organoids (**Fig. 2f, Supplementary Fig. S5a, S5c)**, indicating that this subset remained relatively stable under the current culture conditions. In parallel, naive CD4⁺T cells declined during culture in both groups and were further reduced by day 7 in the stimulated condition (**Fig. 2g, Supplementary Fig. S5a, S5c**), consistent with progressive exit from a naive-like state. By contrast, terminally differentiated (TD) CD4⁺T cells decreased over time (**Fig. 2h, Supplementary Fig. S5a, S5c**). Central memory (CM) CD4⁺T cells, however, increased progressively during culture and reached higher levels by day 7 in both unstimulated and stimulated organoids (**Fig. 2i, Supplementary Fig. S5a, S5c**), suggesting that the culture system promotes time-dependent accumulation of CM-like states, whereas antigen stimulation contributes only modestly to endpoint divergence. CD8⁺T cell subsets showed broadly similar directional changes, including late enrichment of EM-like cells and reduction of naive-like cells, but these trends were weaker overall and more strongly influenced by donor variability (**Supplementary Fig. S5d–g**). Together, these findings indicate that tonsil immune organoids support intrinsic temporal remodeling of T-cell states during culture, while antigen stimulation most clearly accentuates selected adaptive-like features within the CD4⁺T compartment.

We next assessed innate-like lymphoid populations to determine whether the organoid system preserves lymphocyte compartments beyond conventional adaptive B- and T-cell states (**Supplementary Fig. S5b, S5h–j**). CD3⁺CD56⁺ and CD3⁻CD56⁺ populations were both maintained during culture and showed modest temporal fluctuation across days 0, 3, and 7. However, antigen-dependent divergence at day 7 was limited, and these patterns were strongly influenced by donor-to-donor variability. Within the CD3⁻CD56⁺ compartment, CCR7 expression remained broadly high across conditions and time points, suggesting relative preservation of a lymphoid-tissue-associated phenotype rather than a pronounced stimulation-dependent shift. Together, these analyses indicate that tonsil immune organoids retain innate-like lymphoid compartments during culture, but that their remodeling is more limited and less clearly antigen-dependent than the adaptive-like changes observed in B-cell and CD4⁺ T-cell compartments.

Beyond the adaptive-like remodeling observed in B and T cell compartments, we next asked whether antigen-presenting populations also underwent activation-associated changes during culture. In Lin⁻ DC-like cells, HLA-DR median fluorescence intensity increased over time in both unstimulated and antigen-stimulated organoids. Additionally, representative histograms showed a progressive rightward shift of the HLA-DR signal between days 3 and 7 (**Fig. 2j–k**), indicating that antigen-presenting populations undergo progressive activation-associated remodeling during culture. A similar time-dependent increase was observed in CD19⁺ B cells (**Fig. 2l–m**), indicating that B cells not only changed differentiation state during culture but also acquired stronger activation-associated features.

To complement the endpoint and serial imaging analyses, we performed time-lapse imaging of organoids reconstituted with CellTracker 488-labeled T cells and DiD-labeled B cells under IL-21 conditions (**Supplementary Fig. S6**). Images were acquired every 10 min during two observation windows (24–48 h), providing direct dynamic visualization of T- and B-cell redistribution during the emergence of GC-like organization in a M-hTIO setting (*Supplementary Videos 1–2*). These data indicate that our platform sustains coordinated temporal remodeling across both innate and adaptive-like immune dynamics.

## IL-21 and ICOS-L perturbation functionally calibrate GC-like responsiveness in tonsil immune organoids

Having established that tonsil immune organoids support coordinated temporal remodeling across lymphoid and antigen-presenting compartments, we next asked whether this system could resolve functionally distinct GC-helper perturbations. We therefore compared two biologically distinct but complementary perturbations of helper-associated support: IL-21, representing an exogenous downstream helper effector signal, and ICOS-L blockade, representing restriction of an upstream costimulatory axis linked to T–B cell interaction. As illustrated in **Fig. 3a**, IL-21 supplementation and ICOS-L blockade (Prezalumab) are expected to generate distinct phenotype-level response patterns within the organoid system, differing in GC-like timing, B-cell differentiation pace, and downstream B-cell output.

**Figure 3.**
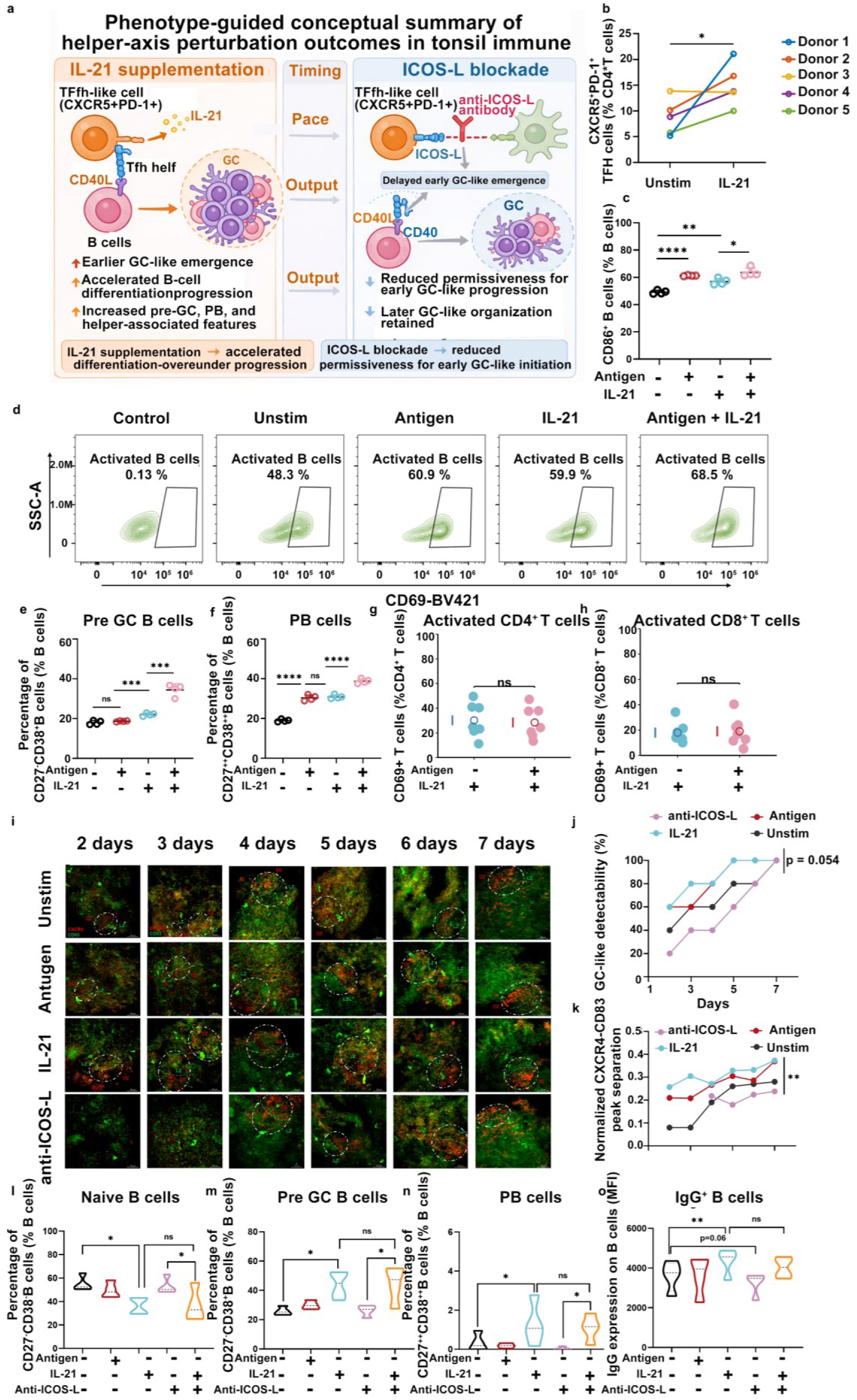
IL-21 accelerates GC-like remodeling and preserves B-cell differentiation under restricted ICOS-L signaling. **a**, Schematic overview of two simplified conceptual frameworks summarizing how IL-21 supplementation and ICOS-L blockade in tonsil immune organoids. The schematics integrate established Tfh–GC biology with the observations obtained in the present study. **b**, Donor-level changes in CXCR5⁺PD-1⁺CD4⁺ Tfh-like cells under unstimulated and IL-21-treated conditions., showing an overall increase in the IL-21 condition across five donors. **c–f,** Quantification of activated B cells (CD86⁺ B cells) (**c-d**), pre-GC B cells (**e**), and plasmablasts (**f**) under the indicated four conditions (unstimulated, NTD, IL-21, and NTD+IL-21) after 7 days of organoid culture. **g–h,** Quantification of CD69⁺ CD4⁺ T cells (**g**) and CD69⁺ CD8⁺ T cells (**h**) in IL-21-containing cultures with or without NTD stimulation at day 7. **i,** Representative serial confocal images showing GC-like structures over time under the unstimulated, NTD, IL-21, and anti-ICOS-L conditions. GC-like regions were identified by combined CD83 and CXCR4 staining. **j**, Quantification of GC-like detectability over time. GC-like detectability was calculated as the percentage of imaged fields/wells containing at least one detectable GC-like structure at each time point. Each field/well was scored as GC-like positive or negative based on the presence of a localized CXCR4-enriched and/or CD83-enriched GC-like region. k, Representative normalized CXCR4–CD83 peak separation in displayed GC-like regions. Peak separation was calculated from line-scan profiles as the distance between dominant CXCR4 and CD83 peaks, normalized to the Feret diameter of the corresponding GC-like ROI. Images without detectable GC-like structures were excluded from this analysis. **l-o**, Frequencies of naive B cells (**l**), pre-GC B cells (**m**), PBs (**n**), and IgG⁺ B cells (**o**) in unstimulated, NTD, IL-21, anti-ICOS-L, and IL-21+anti-ICOS-L groups after 7 days of organoid culture, showing that exogenous IL-21 preserves substantial differentiation output under restricted ICOS-L signaling (n = 4). Data are shown as individual donors, with mean values indicated. Statistical comparisons were performed using unpaired two-tailed t-tests for the indicated comparisons. ns, not significant; *P < 0.05, **P < 0.01, ***P < 0.001, ****P < 0.0001.

We first assessed whether IL-21 could enhance Tfh-associated and B-cell activation phenotypes in this system. In the CXCR5⁺PD-1⁺CD4⁺ compartment, donor-level trajectories showed a consistent increase in the IL-21-treated condition across all five donors (**Fig. 3b**), suggesting that IL-21 supports a Tfh-like response in tonsil immune organoids. At the level of B-cell activation, CD86⁺ B cells were increased in both the NTD and IL-21 groups relative to the unstimulated condition, and the NTD+IL-21 group was further elevated relative to IL-21 alone (**Fig. 3c-d**), indicating that IL-21 can further amplify B-cell activation in the presence of antigen stimulation.

We next examined how IL-21 affected B-cell differentiation. In pre-GC B cells, the IL-21 group was significantly increased relative to the NTD group, and the NTD+IL-21 group was further increased relative to IL-21 alone (**Fig. 3e**), suggesting that IL-21 promotes entry into an early GC-associated differentiation state. A similar pattern was observed for plasmablasts (PBs). IL-21 significantly increased PB frequencies relative to the unstimulated condition, and the NTD+IL-21 group further exceeded IL-21 alone (**Fig. 3f**), indicating synergistic promotion of plasmablast-like output by antigen and IL-21. GC B cells showed only limited separation among groups at this time point (**Supplementary Fig. S7a**). By contrast, naive B cells and memory B cells were both reduced in the IL-21-containing conditions (**Supplementary Fig. S7b-c**), consistent with a shift away from less differentiated B-cell states. Taken together, these results suggest that the dominant effect of IL-21 in this setting is not necessarily expansion of a single terminal GC B-cell compartment at one endpoint, but rather an increase in the differentiation flux driving B cells from activation into pre-GC and plasmablast-like states.

We then examined whether these changes were accompanied by broader alterations in the T-cell compartment. Despite the increase in CXCR5⁺PD-1⁺CD4⁺ Tfh-like cells, the overall proportions of CD4⁺ and CD8⁺ T cells remained comparable across the four conditions (**Supplementary Fig. S7d-g**). Moreover, within the IL-21-containing groups, addition of antigen did not further alter the frequencies of CD69⁺T cells in either the CD4⁺ or CD8⁺ T cell compartments at day 7 (**Fig. 3g-h**). These observations suggest that, at this endpoint, IL-21 does not primarily act by broadly increasing terminal T-cell activation, but rather by promoting a more selective helper-associated program that is coupled to enhanced B-cell differentiation output.

We next examined how GC-helper dynamics changed when ICOS-L signaling was restricted. Because ICOS-L is linked to helper-dependent GC organization, we first assessed GC-like emergence by serial confocal imaging (**Fig. 3i-k)**. In this system, GC-like structures first became detectable on day 2 and matured progressively from day 3 to day 5. Quantification of GC-like-positive fields showed that IL-21-treated organoids showed earlier appearance of clearly detectable GC-like regions, consistent with accelerated GC-like emergence. By contrast, in the anti-ICOS-L condition, GC-like structures were less readily detectable during the early phase, but became visible at later time points and were clearly present by day 7 (**Fig. 3j)**. In parallel, quantification of DZ-like/LZ-like zonation showed that clearer separation between CXCR4-enriched and CD83-enriched domains was associated with more evident GC-like organization. In this framework, greater zonation clarity was interpreted as clearer GC-like spatial organization, reflecting more pronounced segregation between DZ-like and LZ-like regions (**Fig. 3k)**. These observations suggest that ICOS-L restriction primarily delays the onset of GC-like formation rather than preventing its later development.

We then asked whether this kinetic delay was reflected in day-7 phenotypes. Across the five-group comparison (unstimulated, NTD, IL-21, anti-ICOS-L, and IL-21+anti-ICOS-L). Relative to the unstimulated condition, anti-ICOS-L did not cause complete loss of later GC-like structures or broad collapse of all downstream phenotypes. However, the B-cell compartment under anti-ICOS-L retained a comparatively less differentiated profile. Naive B cells remained higher, whereas pre-GC B cells and plasmablasts showed lower tendencies than in more strongly differentiated conditions. Naive B cells remained comparatively higher, whereas pre-GC B cells and plasmablasts were lower than in more strongly differentiated conditions (**Fig. 3l-m)**. Consistent with this pattern, IgG^+^ B cell signal over the unstimulated condition, and both anti-ICOS-L and IL-21+anti-ICOS-L tended to remain lower than IL-21 alone at this endpoint **(Fig. 3o)**. GC B and memory B cells showed little separation across groups at this endpoint (**Supplementary Fig. S7h-i).** Together, these findings suggest that ICOS-L restriction in this system primarily delays early GC-like initiation and limits progression toward more differentiated B-cell states, rather than causing complete loss of later GC-like structures. Notably, addition of exogenous IL-21 largely preserved the downstream differentiation program under ICOS-L restriction. although some readouts, including IgG-associated signal, still trended lower than with IL-21 alone, indicating that exogenous IL-21 was sufficient to maintain much of the differentiation-associated program even when the ICOS-L axis was constrained.

These findings indicate that the organoid system does not merely undergo passive culture-associated drift, but remains responsive to defined helper-axis inputs and can be directionally modulated in a biologically interpretable manner, thereby providing a functional reference for the adjuvant studies presented in the following section.

## Pre-exist Memory Shapes Adjuvant Gene Profile Responses

Because vaccine responses occur in the context of prior antigen exposure, we asked whether distinct recall-conditioned immune baselines would alter the transcriptional programs induced by adjuvant stimulation. To address this, we designed three pre-existing immune models, representing progressively different antigen-experience states, with M-hTIO framework.

In this workflow (**Fig. 4a**), organoids generated from four COVID-19 convalescent donors were organized into three experimentally defined memory contexts. We designed three cohorts, representing different baseline immune states, according to donors’ prior antigen exposure history: Adjuvant-prime Organoids (AO) comprised organoids primed with HA at day 0 and subjected to adjuvant-only secondary stimulation at day 7. Weak pre-existing immune Organoids (WPO) comprised organoids primed with HA and re-stimulated on day 7 with blank, HA alone, or HA-formulated vaccine, representing a weak pre-existing immune memory setting. Strong Pre-existing Organoids (SPO) comprised organoids conditioned with RBD, and were re-stimulated on day 7 with blank, RBD alone, or RBD-formulated vaccine, representing a strong pre-existing immune memory setting. These operational definitions allowed us to compare adjuvant responses under no-memory, weak-memory, and strong-memory organoid settings within the same M-hTIO framework. However, although clinically tested adjuvants have established dose ranges in vivo and in conventional cell systems, we were unable to identify directly transferable reference concentrations for their use in a human immune organoid context. We therefore first performed an organoid-based benchmarking step to define working concentrations for five clinically tested adjuvants together with two antigens. To do this, each condition was evaluated empirically after 24 h using two complementary criteria, including preservation of viability and induction of early immune activation. Serial concentrations of HA, RBD, Al, Mnβ, ODN1018, R848, and AddaSO_3_ were tested, and concentration-dependent trends were jointly evaluated to define organoid-compatible response windows (**Supplementary Fig. S8–S10**). This analysis identified organoid-compatible concentration windows in which tissue integrity was retained while measurable immunostimulatory activity remained detectable, thereby avoiding conditions that were either poorly active or overly disruptive to the organoid system. After 24 h of stimulation, bulk RNA-seq revealed clear transcriptional segregation among tonsil organoid models with distinct pre-existing immune baselines, with SPOs clustering separately from AOs and WPOs (**Fig. 4b**). Differential expression analysis of the blank SPO and WPO groups identified marked baseline divergence: SPOs were enriched for B cell and antigen-presentation genes (*MS4A1*, *CD79B*, *B2M*, and *CD74*), whereas WPOs preferentially expressed genes linked to innate immunity, complement, and stress adaptation (*C1QB*, *FAIM2*, *RASD1*, and *BPIFB4*) (**Fig. 4c**). Consistently, KEGG analysis showed enrichment of microenvironment- and signaling-related pathways in WPOs, including ECM-receptor interaction, focal adhesion, and calcium/cyclic nucleotide signaling, while SPOs were enriched for canonical immune activation pathways, including antigen processing and presentation, B cell receptor signaling, NF-κB signaling, Fc gamma receptor-mediated phagocytosis, and T helper cell differentiation. These findings indicate that weak and strong pre-existing immune models are distinguished by innate microenvironmental regulation versus adaptive immune activation, respectively.

**Figure 4.**
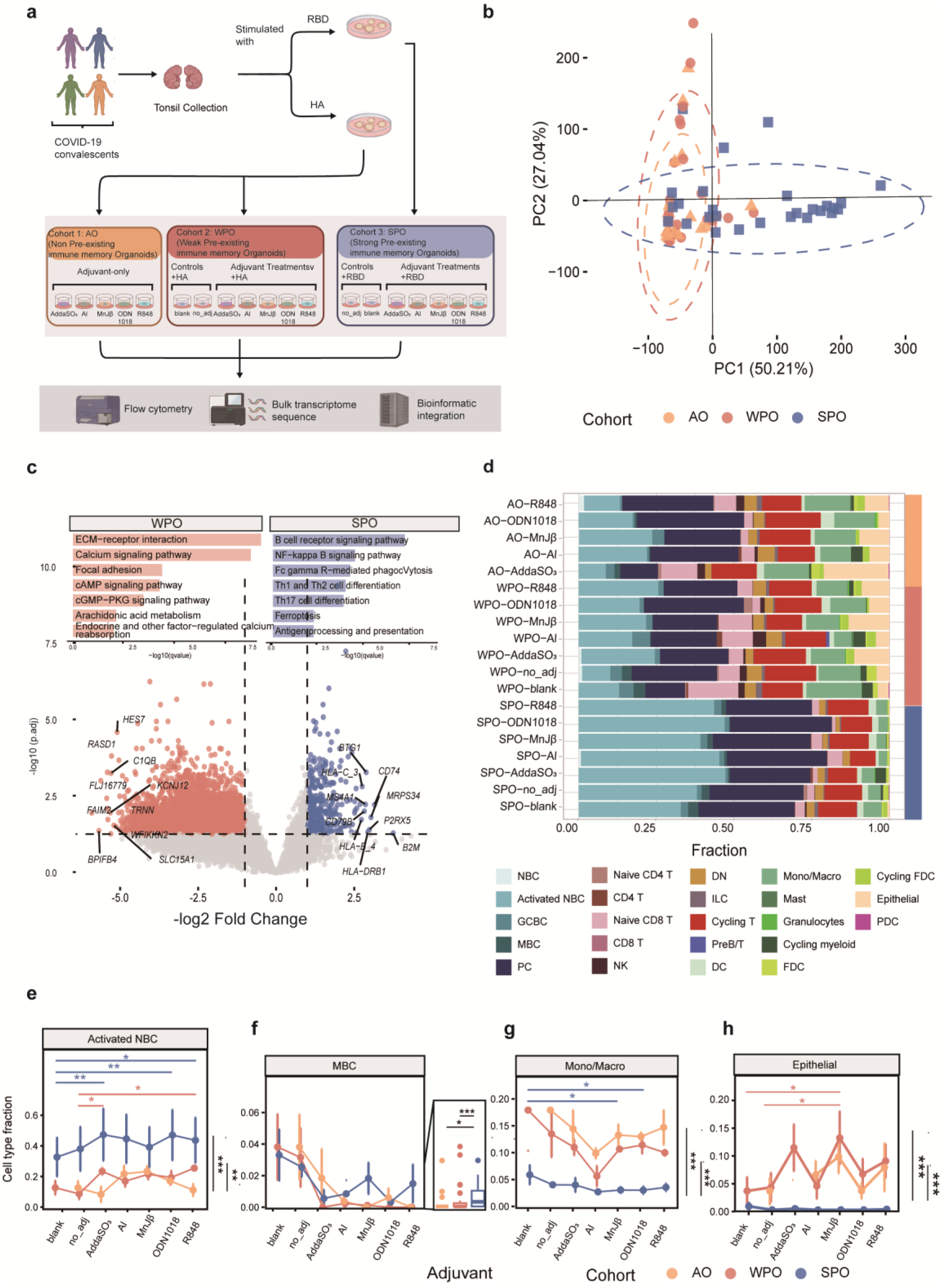
Pre-existing immune level reshape organoid’s adjuvant gene responses. **a**, Work flow of generation and stimulation of different pre-existing immune baseline M-hTIO. **b,** PCA result of bulk transcriptome sequence data after 24h’s stimulation. Cohort types are labeled with different color; **c,** Volcano plot of bulk RNA expression data between SPO and WPO, GO enrichment analysis of DEGs are shown in the bar plot. **d,** Deconvolution of bulk transcriptomic data. **e-g**, Significantly enriched cell type’s fraction in each cohort. Significance among SPO, WPO and AO are shown in the right panel, using wilcox test and adjusted with BH method, the panel on top of the lines is the significance among different groups between each adjuvant group to their no_adj or blank control, using Linear Mixed-Effects Model (LMM) and post-hoc pairwise comparisons between each adjuvant group and the blank control. P-values were adjusted for multiple comparisons using Dunnett’s method. *p < 0.05, **p < 0.01, ***p < 0.001

Although antigen alone induced limited additional divergence, adjuvant inclusion further enhanced model-specific immune responses, demonstrating the utility of this platform for assessing antigen-adjuvant effects **(Supplementary Fig. S11)**. To characterize cellular microenvironmental changes induced by different vaccine formulations, we performed deconvolution analysis of bulk transcriptomic data (**Fig. 4d**). Pre-existing immune status emerged as the major source of variation across cohorts. SPOs exhibited a markedly lymphoid-enriched microenvironment, with prominent enrichment of B cell populations, particularly activated NBCs and MBCs (**Fig. 4e, f**). In contrast, WPOs and AOs showed higher baseline proportions of monocytes/macrophages and epithelial cell, accompanied by the absence of the strong humoral signature observed in SPOs (**Fig. 4g, h**). In addition, adjuvant treatment further reshaped cellular composition within each model (**Fig. 4e-h**), consistent with distinct immunomodulatory effects of different formulations. Together, these inferred cellular proportions indicate that the local immune landscape is primarily determined by antigen exposure history, with adjuvants providing a secondary layer of modulation.

To assess pathway-level dynamics across stimulation conditions, we performed Gene Set Variation Analysis (GSVA) on RNA-seq data and focused on pathways significantly upregulated in at least one adjuvant-treated group relative to the matched no-adjuvant control. Unsupervised clustering separated SPOs from AOs and WPOs, with the latter two showing similar globel adjuvant-responsive signatures (**Fig. 5a**). AOs and WPOs were enriched for pathways related to tissue repair, inflammatory remodeling, innate immune infiltration, and metabolic adaptation (**Fig. 5a, cluster 1**), whereas SPOs preferentially enriched adaptive recall-associated programs, including B cell receptor signaling, T cell activation/proliferation, antigen presentation, cytokine responses, and Fc gamma receptor signaling (**Fig, 5a, cluster 2**).

**Figure 5.**
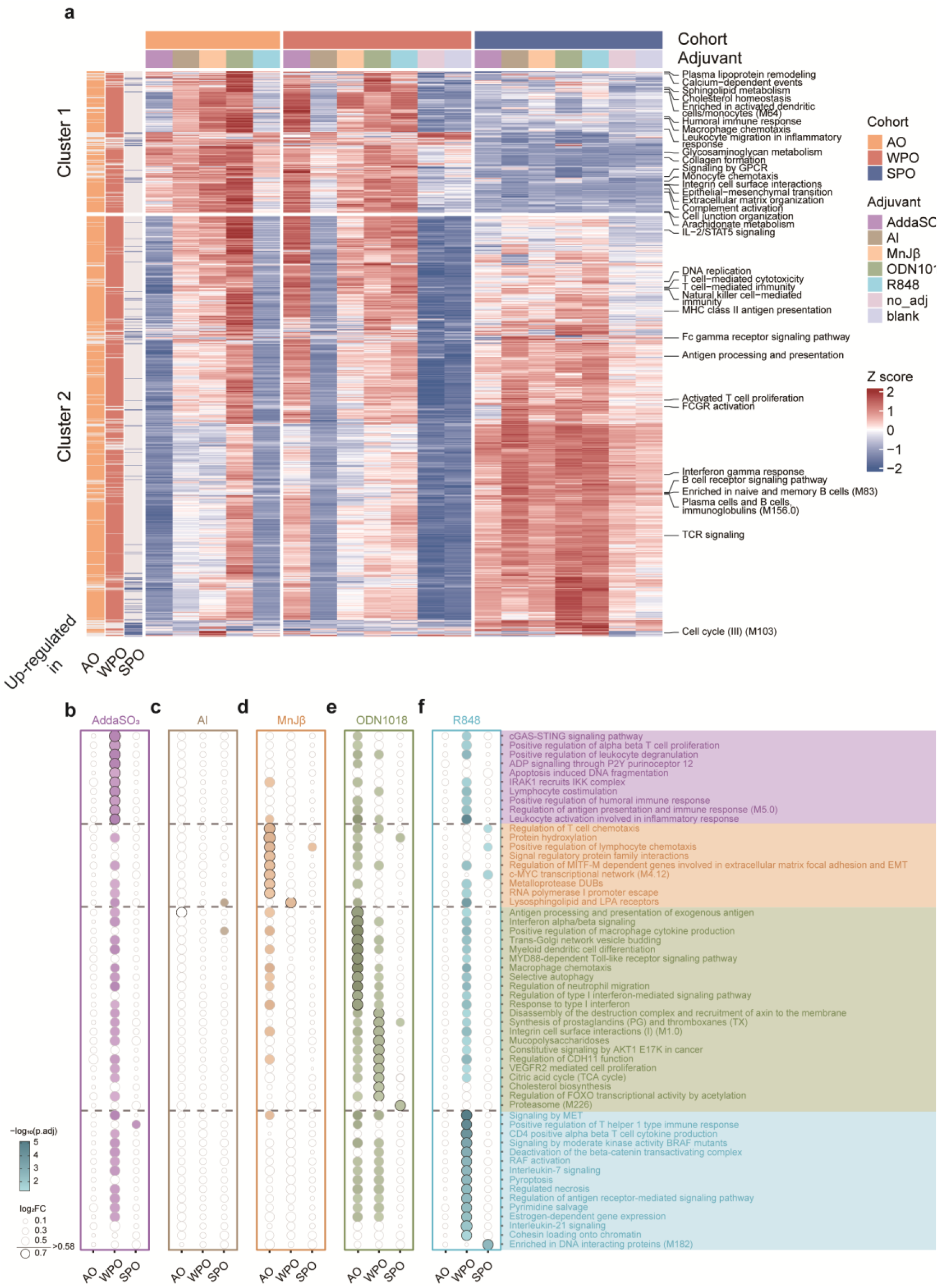
GSVA reveals divergent mechanisms of adjuvant-induced immune pattern among models. **a**, Overall pathway activation score heatmap. Pathways that significantly upregulated in at least one adjuvant group compared to no_adj control within each cohorts. Group information is showen in the top panel. Scores are normalized as in the plot. Pathways were unsupervised clustering into two parts with typical pathways labeled on the right. **b,** Specific upregulated pathways for each group. Pathways exhibiting statistical significance (FDR < 0.05 for WPO/AO cohorts, p < 0.05 for SPO cohorts) are highlighted in color, where circle size indicating the expression fold change while those with log2(Fold change) > 0.58 are marked with black circles.

We further evaluated the similarity of adjuvant responses within each model (**Supplementary Fig. S12a**). In AOs and WPOs, different adjuvants displayed more heterogeneous response patterns, while SPOs were highly concordant, likely reflecting dominant recall of pre-existing immune memory that masked additional adjuvant-specific effects.

Together, these findings highlight the utility of baseline-stratified organoid models for dissecting early adjuvant responsiveness and demonstrate that pre-existing immune state is a key determinant of vaccine-induced transcriptional programs.

## Transcriptomic profiling reveals divergent and shared mechanisms of adjuvant-induced immune activation

We further investigated the immune functions of different adjuvants during the early phase of vaccine responses. Based on the GSVA results, we identified differentially enriched pathways associated with each adjuvant (**Fig. 5b–f**).

For AddaSO_3_ (**Fig. 5b**), functional enrichment analysis revealed a coordinated immune program progressing from DAMP-associated innate inflammatory sensing to adaptive immune activation. Enriched innate pathways included apoptosis-induced DNA fragmentation, cGAS–STING signaling, ADP signaling through P2Y purinoceptor 12, and leukocyte activation in inflammatory responses. Pathways involved in antigen presentation, lymphocyte costimulation, alpha-beta T cell activation, and humoral immune responses were also upregulated, indicating coordinated innate and adaptive immune activation.

Al adjuvant failed to induce significant pathway enrichment within the early time examined (**Fig. 5c**), consistent with previous reports that alum generally exhibits relatively limited early inflammatory and cell-mediated immunostimulatory activity compared with more potent innate immune agonists^[16]^.

MnJβ induced a transcriptomic program marked by lymphocyte recruitment, metal-dependent enzymatic activity, and transcription factor-driven proliferative expansion (**Fig. 5d**). This response was characterized by enrichment of pathways associated with T cell and lymphocyte chemotaxis, together with MITF-M-associated networks linked to extracellular matrix remodeling, focal adhesion, and EMT-like processes. Increased SIRP family signaling pointed to enhanced receptor–ligand crosstalk between myeloid antigen-presenting cells and lymphocytes. In parallel, MnJβ upregulated the c-Myc transcriptional network together with RNA polymerase I promoter escape pathways, consistent with increased biosynthetic activity. MnJβ also showed significant upregulation of lysosphingolipid and lysophosphatidic acid receptor pathways upon additional antigen encounter.

For ODN1018, the transcriptional response revealed a coordinated program linking innate immune sensing to adaptive immune activation and tissue microenvironment remodeling (**Fig. 5e**). ODN1018 induced a canonical MyD88-dependent innate immune cascade characterized by interferon-α/β and Toll-like receptor signaling, together with chemotactic programs involving macrophages, neutrophils, and dendritic cells. Pathways related to vesicular trafficking, Golgi biogenesis, and selective autophagy were also upregulated, consistent with remodeling of the endolysosomal system to support receptor processing and MHC class II-mediated antigen presentation. Adaptive immune pathways associated with CD4^+^ /CD8^+^ T cell differentiation, B cell affinity maturation, and humoral immunity were concomitantly enriched. In parallel, ODN1018 promoted metabolic and stromal remodeling, as reflected by enrichment of the tricarboxylic acid cycle, cholesterol biosynthesis, PI3K/AKT/FOXO signaling, prostaglandin synthesis, VEGFR2-mediated angiogenesis, and integrin- or cadherin-11-dependent adhesion pathways.

R848 elicited a transcriptional program indicative of viral sensing mimicry and adaptive immune activation (**Fig. 5f**). Enriched pathways included positive regulation of Th1-type immune responses and CD4^+^ αβ T cell cytokine production, consistent with induction of Th1-skewed cellular immunity. Upregulation of interleukin-21 signaling suggested support for follicular helper T cell function, B cell affinity maturation, and immunoglobulin class switching, whereas enrichment of interleukin-7 signaling was consistent with enhanced survival of effector T cells. Increased antigen receptor-mediated signaling further indicated amplification of both TCR- and BCR-dependent activation programs.

Furthermore, the pathway enrichments identified align closely with established immunological paradigms (**Supplementary Fig. S12b**). For instance, the observed transcriptional quiescence of traditional aluminum salts, TLR9-mediated innate priming by ODN1018 and the TLR7/8-driven adaptive Th1 skewing by R848 are well-documented phenomena in adjuvant research^[17–20]^.

In summary, the 24-hour comparative transcriptomic landscape revealed distinct immune dynamics: AddaSO_3_ promoted coordinated innate inflammatory sensing and adaptive activation, alum remained transcriptionally quiescent, MnJβ was associated with lymphocyte recruitment and biosynthetic activity, ODN1018 drove TLR9/MyD88-dependent innate priming together with metabolic and stromal remodeling, and R848 preferentially induced a viral sensing-like, Th1-skewed adaptive program.

Because adjuvant-induced responses varied across antigen settings, with some effects emerging more prominently in specific baseline contexts, we next sought to distinguish context-dependent responses from transcriptional features that were consistently attributable to each adjuvant. To this end, we modeled the main effect of adjuvant treatment while accounting for inter-individual baseline variation and antigen-specific non-additive effects (**Fig. 6a–d**). Distinct adjuvant-specific transcriptomic signatures emerged from this analysis. AddaSO_3_ preferentially induced stress- and remodeling-associated genes, including HMOX1, MMP14, MMP1, CTSD, and S100A9 (**Fig. 6a**). MnJβ was characterized by increased expression of KRT and SPRR family members, consistent with an epithelial-associated transcriptional program (Fig. 6b). Among the tested conditions, ODN1018 elicited the broadest transcriptional shift, with widespread transcript downregulation accompanied by induction of inflammatory and antigen-presentation-related genes such as CXCL8, HLA-DRB1, LYZ, and CTSD (**Fig. 6c**). In contrast, R848 induced a comparatively restricted response marked by NFKBIA and a limited set of inflammatory genes (Fig. 6d). No significant conserved transcriptional signature was observed for Al.

**Fig. 6.**
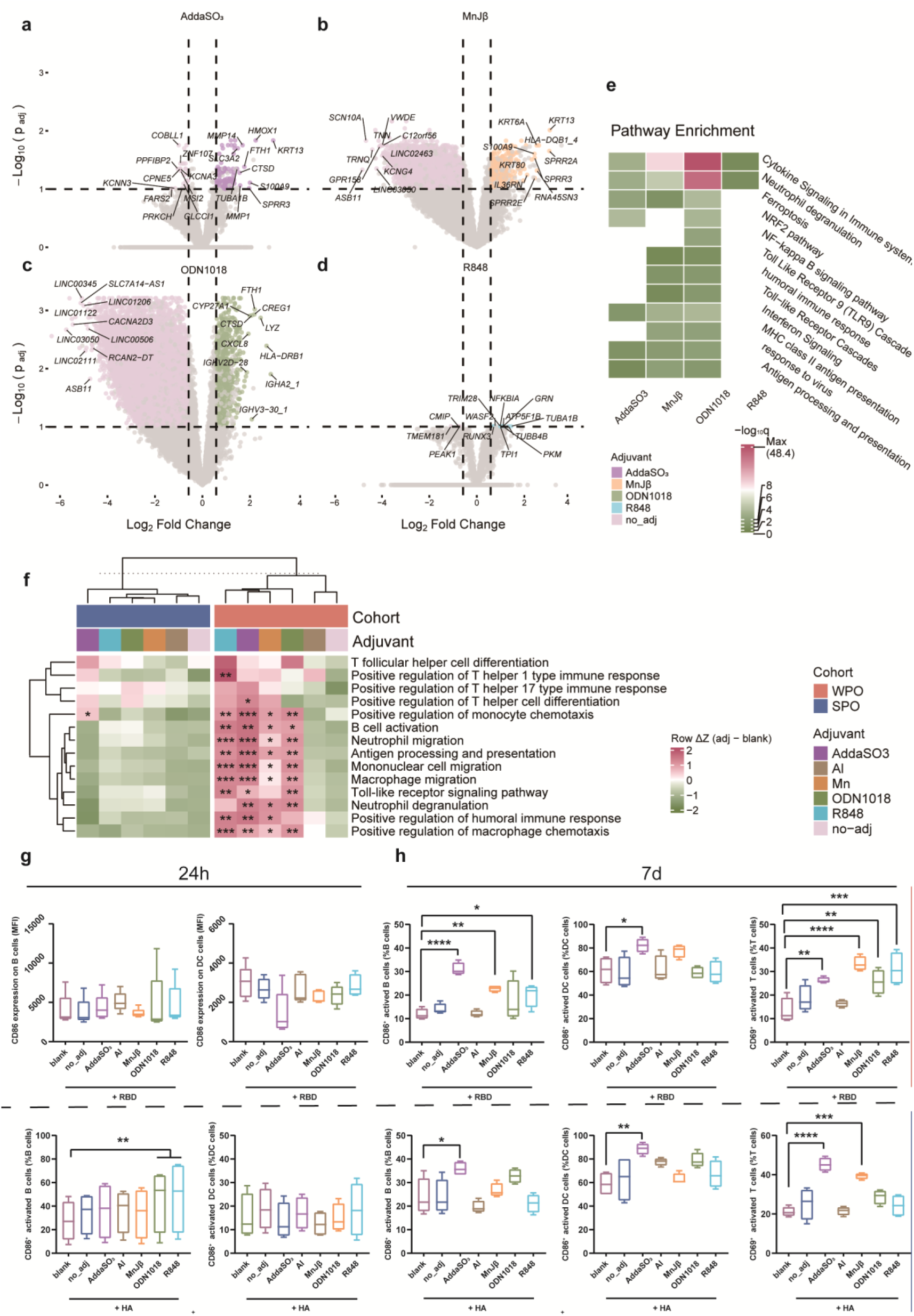
Transcriptomic Profiling Reveals Divergent and Shared Mechanisms of Adjuvant-Induced Immune Activation. **a-d**, Conserved gene volcano plots for AddaSO_3_, MnJ β , ODN1018, R848,with cutoff as p.adj < 0.1, log_2_ FC > 0.585. **e,** GO enrichment analysis for conserved gene sets. **f,** Heatmap of vaccine function related pathways GSVA enrichment score. Colors are shown as the difference between each group to their relative blank control. P values are labeled when p.adj between adj group and blank control are less than 0.05. **g-h,** Activated Cell evaluation for each group after 24h or 7d. n = 4 independent donors per group. Statistical significance compared to the blank group was determined using Repeated Measures (RM) one-way ANOVA followed by Dunnett’s multiple comparisons test. *P < 0.05, **P < 0.01.

Functional enrichment analysis of upregulated genes revealed marked differences in pathway usage across adjuvants (**Fig. 6e**). AddaSO_3_ enriched cytokine signaling, neutrophil degranulation, ferroptosis, and NRF2 signaling pathways, whereas MnJβ was dominated by interferon signaling, antiviral response, MHC class II antigen presentation, and cytokine signaling. ODN1018 preferentially engaged TLR9/Toll-like receptor cascades, NF-κB signaling, antigen processing and presentation, and neutrophil degranulation, while R848 displayed a narrower profile centered on cytokine signaling and neutrophil degranulation. Together, these results reveal distinct functional programs associated with each adjuvant. Protein–protein interaction analysis of the core response genes identified multiple densely connected modules enriched for peptide chain elongation, neutrophil degranulation, cytokine signaling, apoptotic regulation, and endoplasmic reticulum stress (**Supplementary Fig. S13a**). These modules showed substantial gene overlap among AddaSO_3_, MnJβ, and ODN1018. Consistently, transcription factor enrichment analysis revealed a shared regulatory core across these three adjuvants, centered on NFKB1, RELA, STAT3, and CEBPB, with ODN1018 showing the strongest enrichment (**Supplementary Fig. S13b, c**). Adjuvant-specific features were also evident, including IRF1/BCL6 programs in ODN1018, ELF3-linked targets in MnJβ, and CREB1-associated targets in AddaSO_3_. Additional enrichment of HIF1A/MYC-, XBP1/ATF4/ATF6-, and PPARA/PPARG-associated programs suggested engagement of metabolic, ER stress, and lipid-related regulatory pathways. By contrast, R848 showed a more distinct pattern with weaker representation of this shared network. Together, these findings indicate that AddaSO_3_, MnJβ, and ODN1018 converge on partially shared functional and regulatory hubs, whereas R848 engages a more distinct transcriptional program.

Having defined both the distinct and shared transcriptional features of individual adjuvants, we next asked how these programs were manifested in vaccine settings with different pre-existing immune baselines. GSVA of curated vaccine-relevant pathways revealed a clear dependence of adjuvant activity on the baseline immunogenic context of the antigen (**Fig. 6f**). Overall, pathway induction was stronger in WPOs than in SPOs. In the weak pre-existing immune setting, R848, AddaSO_3_, ODN1018, and Mn each enhanced multiple immune modules, with R848 and AddaSO_3_ showing the broadest enrichment across innate cell recruitment, Toll-like receptor signaling, antigen presentation, B cell activation, and humoral immune pathways. In contrast, AddaSO_3_ was the dominant inducer in the strong pre-existing immune antigen group, where enrichment was largely restricted to cell migration and innate immune programs.

Further, we assessed immune cell composition and activation states across antigen–adjuvant conditions at 24 h and day 7. At 24 h, significant changes were limited, with increased frequencies of CD86^+^ activated B cells observed only in the WPO group treated with R848 or ODN1018. By day 7, broader adjuvant-associated effects emerged in both the SPO and WPO groups. AddaSO_3_ induced the most extensive long-term changes, significantly increasing the proportions of CD86^+^ activated B cells, CD86^+^ dendritic cells, and CD69^+^ activated T cells in both cohorts. MnJβ was likewise associated with a strong day-7 response, most prominently in activated T-cell populations. By contrast, the B-cell activation seen with R848 and ODN1018 was diminished at day 7.

Collectively, these analyses reveal that adjuvants induce both distinct and partially convergent immune programs across early and sustained phases of vaccine responses. Although each adjuvant exhibited a characteristic transcriptional and functional profile, ODN1018, AddaSO_3_, and Mn converged on NF-κB/STAT3-associated regulatory modules, while cellular analyses identified more pronounced differences at later time points. Together, these findings indicate that adjuvant activity reflects both adjuvant-intrinsic regulatory programs and context-dependent shaping by antigen baseline immunogenicity.

Together, these data establish a multi-layered view of adjuvant action during vaccine stimulation. Early transcriptomic profiling identified adjuvant-specific pathway signatures, with AddaSO_3_, MnJβ, ODN1018, and R848 each engaging different combinations of innate, adaptive, and tissue-remodeling pathways, whereas alum remained largely quiescent. Core-signature modeling further resolved distinct adjuvant-intrinsic transcriptional programs and pathway architectures, while transcription factor analysis identified convergence of ODN1018, AddaSO_3_, and Mn on NF-κB/STAT3-centered regulatory modules. Adjuvant effects were further shaped by antigen baseline immunogenicity, with stronger pathway induction in weak pre-existing immune settings.

## Discussion

Human vaccine responses are shaped by the interplay between antigenic context, pre-existing immune state, and adjuvant-driven immune programming. In this study, we established a low matrix supported human tonsil immune platform, M-hTIO, and applied it to comparative adjuvant evaluation under distinct recall-conditioned immune baselines. By integrating GC-like morphological remodeling, dynamic T–B cell interactions, early transcriptional profiling, and longer-term adaptive immune phenotyping, this system allowed us to examine vaccine responsiveness across multiple temporal and functional layers. Rather than ranking adjuvants solely by the magnitude of acute activation, our platform distinguishes baseline immune memory, early adjuvant-driven stimulation, and sustained adaptive remodeling. This distinction is particularly relevant for human vaccinology, where prior antigen exposure, vaccination history, infection background, and donor-to-donor heterogeneity can substantially influence subsequent immune responses.

A key finding of this study is that early immune activation and prolonged adaptive remodeling are not always concordant. The 24-h stimulation window captured rapid transcriptional and phenotypic changes induced by adjuvant-containing formulations, whereas extended culture revealed whether these early responses could be maintained and converted into sustained B cell-, DC-, and GC-like remodeling. This temporal distinction is important because a strong early activation signal does not necessarily indicate a productive adaptive response. Therefore, temporally resolved evaluation may provide a more informative framework for adjuvant selection than single-time-point measurements.

The antigen-conditioned design further allowed us to assess how immune baseline states influence adjuvant responsiveness. Human vaccine responses rarely occur in a fully naïve immune environment; prior infection, vaccination, and tissue-resident memory can shape subsequent responses to antigen–adjuvant stimulation. By comparing adjuvant-only, weak-memory, and strong-memory backgrounds, M-hTIO provides a strategy to distinguish adjuvant-driven activation from antigen-associated memory effects. This feature may be particularly useful for evaluating vaccine formulations in human populations with heterogeneous immune histories.

Among the tested adjuvants, AddaSO_3_ showed a relatively consistent activity pattern across early and prolonged response phases, supporting both rapid immune activation and sustained adaptive remodeling. In contrast, some adjuvants induced strong early signatures but showed less consistent prolonged effects, highlighting the importance of evaluating immune trajectory rather than acute potency alone. These findings suggest that productive adjuvant activity should be defined by a balance between early immune activation, cellular viability, B cell and DC remodeling, and maintenance of GC-like organization.

An interesting feature of the helper-axis calibration experiments was that anti-ICOS-L alone produced only limited separation from the unstimulated condition in several day-7 flow-cytometric readouts, while serial imaging suggested delayed early GC-like emergence rather than failure of later formation. Considered together with the similarly modest separation between IL-21 and IL-21 plus anti-ICOS-L, these findings raise the possibility that ICOS–ICOSL signaling in this system contributes mainly to early Tfh-associated organization and GC-like initiation, whereas exogenous IL-21, acting as a more downstream helper effector, may partially compensate for upstream ICOS-L restriction and preserve much of the subsequent B-cell differentiation program. This interpretation is broadly consistent with published work implicating the ICOS–ICOSL axis in Tfh differentiation and T–B coordination, and IL-21 in downstream GC-associated progression and plasmablast differentiation^[21]^. Alternatively, ICOSL blockade may induce compensatory pathways that partially maintain organoid homeostasis despite impaired canonical T–B cooperation. Identifying these alternative mechanisms, and determining whether they involve CD40–CD40L signaling, cytokine compensation, altered antigen presentation, or innate immune rewiring, will be an important direction for future study.

Several limitations should be acknowledged. First, although M-hTIO recapitulates key GC-like features, it remains an ex vivo model and cannot fully reproduce the anatomical, vascular, stromal, and migratory complexity of lymphoid organs in vivo. Therefore, GC-like structures observed in this study should be interpreted as organized adaptive immune remodeling rather than fully mature germinal centers. Second, donor heterogeneity is both a strength and a limitation. While human tonsil samples preserve physiologically relevant immune diversity, differences in infection history, vaccination background, tissue inflammation, and baseline memory may influence adjuvant responsiveness and reduce statistical power. Third, the current study mainly relies on transcriptional, phenotypic, and imaging-based readouts. Future studies incorporating antigen-specific B cell tracking, BCR/TCR sequencing, spatial transcriptomics, antibody affinity measurement, and functional neutralization assays will be needed to further validate the adaptive quality of the responses. Finally, different adjuvants may have distinct optimal dose windows in organoid culture, and strong innate agonists may induce early activation while compromising long-term viability or tissue-like organization. Systematic dose–response and time-course studies will therefore be important for future optimization.

Together, our study establishes M-hTIO as a human-relevant platform for systematic and comparative adjuvant evaluation. By distinguishing rapid immune activation from sustained adaptive remodeling across different immune memory baselines, this platform provides a framework for rational adjuvant selection and high-throughput systems vaccinology.

## Resource availability

### Lead contact

Further information and requests for resources, reagents, or additional details related to this protocol should be directed to and will be fulfilled by the lead contact, Xiao Liu.

### Technical contact

Technical questions or requests for guidance on executing this protocol, including tissue dissociation, organoid culture, and immunostaining procedures, should be directed to and will be answered by Dandan Meng.

### Materials availability statement

- This study did not generate new unique reagents or cell lines.
- All human tonsil samples were obtained from consenting donors in accordance with institutional ethical guidelines.
- All commercially available reagents and materials used in this study, including organoid culture media and supplements, are listed in the key resources table. Researchers interested in obtaining specific components should refer to their respective suppliers.

### Data and code availability statement

Bulk RNA-seq data generated in this study will be deposited in a public repository before publication. All other data supporting the findings of this study are available within the article and its Supplementary Information files, or from the corresponding author upon reasonable request. Custom scripts used for data processing and visualization are available from the corresponding author upon reasonable request.

## Acknowledgments

This work was supported by funding from the Shenzhen Science and Technology Program [WDZC20220819134430002], the Scientifc Research Start-up Funds [QD2021005N], Department of Chemical Engineering-iBHE Special CooperationJoint Fund [DCE-iBHE-2025-2].

We thank all members of the Institute of Biopharmaceutical and Health Engineering (iBHE), Tsinghua Shenzhen International Graduate School, for their valuable discussions and technical assistance. We also acknowledge the support of the Tsinghua University Core Facilities for Life Sciences for providing access to flow cytometry and confocal microscopy platforms. The graphical abstract was created with BioRender.com.

## Author contributions

**Dandan Meng:** Conceptualization, Investigation, Data curation, Formal analysis, Visualization, Writing – original draft. **Xuqi Li:** Data curation, Visualization, Investigation, Writing – original draft. **Xiuli Rao:** Resources, Investigation. **Xiaofen Huang:** Conceptualization, Methodology. **Qiyu Deng:** Investigation, Validation. **Han Ding:** Investigation, Formal analysis. **Lin Li:** Investigation. **Wentao Ma:** Investigation. **Yuan Tao:** Resources, Project administration. **Xiaohua Feng:** Resources, Project administration. **Xiao Liu:** Supervision, Funding acquisition, Project administration, Writing – review & editing.

