## Supplementary Information for "Harnessing Human Immune Organoid model for Systemic and Comparative Analysis of Clinically Vaccine Adjuvants"

### Supplement File

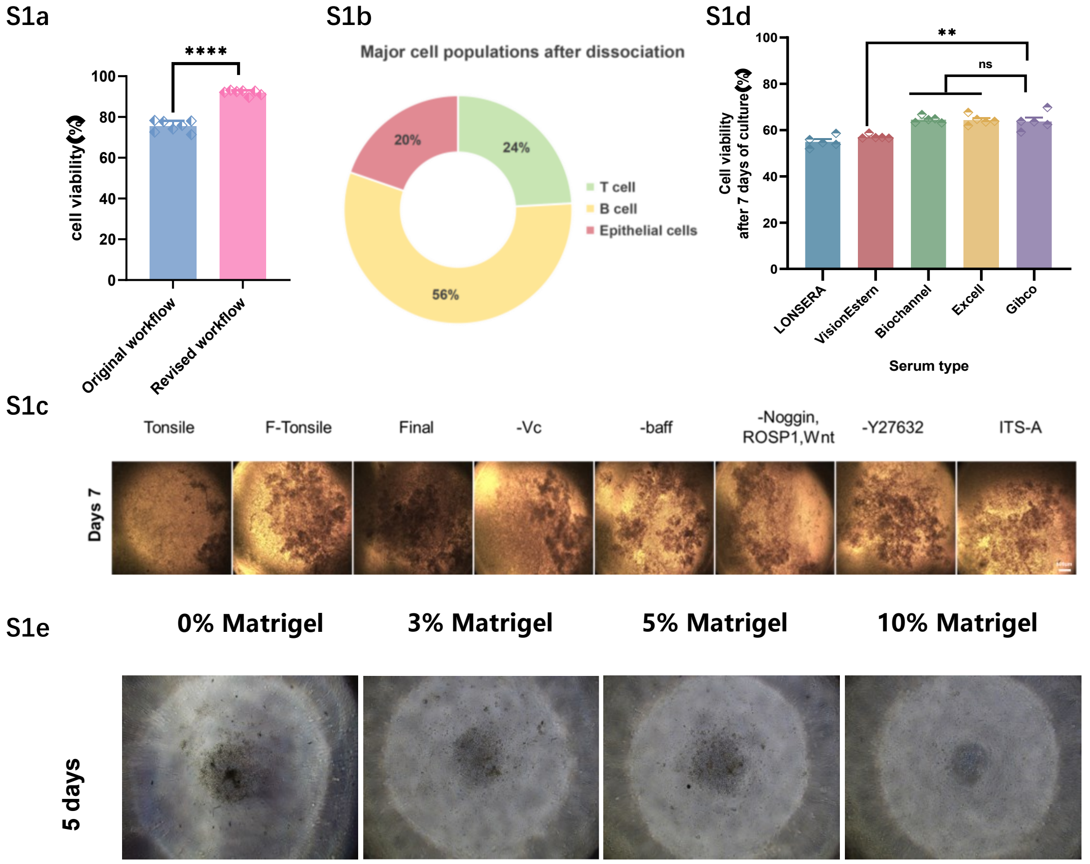


**Supplementary Fig. S1 | Definition of tissue-processing and culture conditions for the human tonsil immune organoid platform. a,** Viability of dissociated tonsil cells obtained with the final tissue-processing workflow compared with the previous procedure. **b,** Flow cytometric analysis of the major cellular composition of dissociated tonsil samples after processing, showing preservation of a lymphocyte-dominant population together with a measurable epithelial fraction. **c,** Representative bright-field images comparing organoid morphology under different culture media after 7 days of transwell culture. Final and F-Tonsile supported the formation of more organoid-like aggregates than Tonsile. **d,** Comparison of commercially available serum sources for tonsil organoid culture. Gibco showed better overall viability, whereas Biochannel and ExCell produced similar results under the tested conditions. **e,** Evaluation of low-concentration Matrigel conditions for transwell culture. Low matrix concentrations, particularly 3%–5%, better preserved aggregate integrity than higher matrix concentrations, and 5% Matrigel was selected for subsequent experiments.

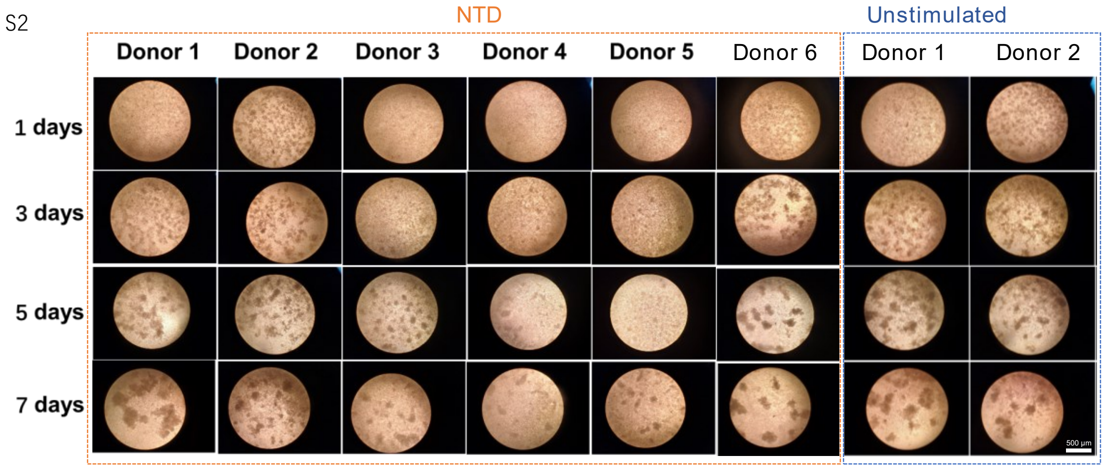


**Supplementary Fig. S2 | Bright-field images of tonsil immune organoid re-aggregation across donors under unstimulated and antigen-stimulated conditions.**

Representative bright-field images showing progressive re-aggregation of dissociated tonsil cells into compact organoid-like clusters at days 1, 3, 5, and 7 under the final culture conditions. Images from multiple donors are shown for both unstimulated and antigen-stimulated cultures. Gross aggregate morphology became progressively more compact over time, whereas no obvious difference was observed between unstimulated and antigen-stimulated groups at the bright-field level. Bar scale = 500 μm

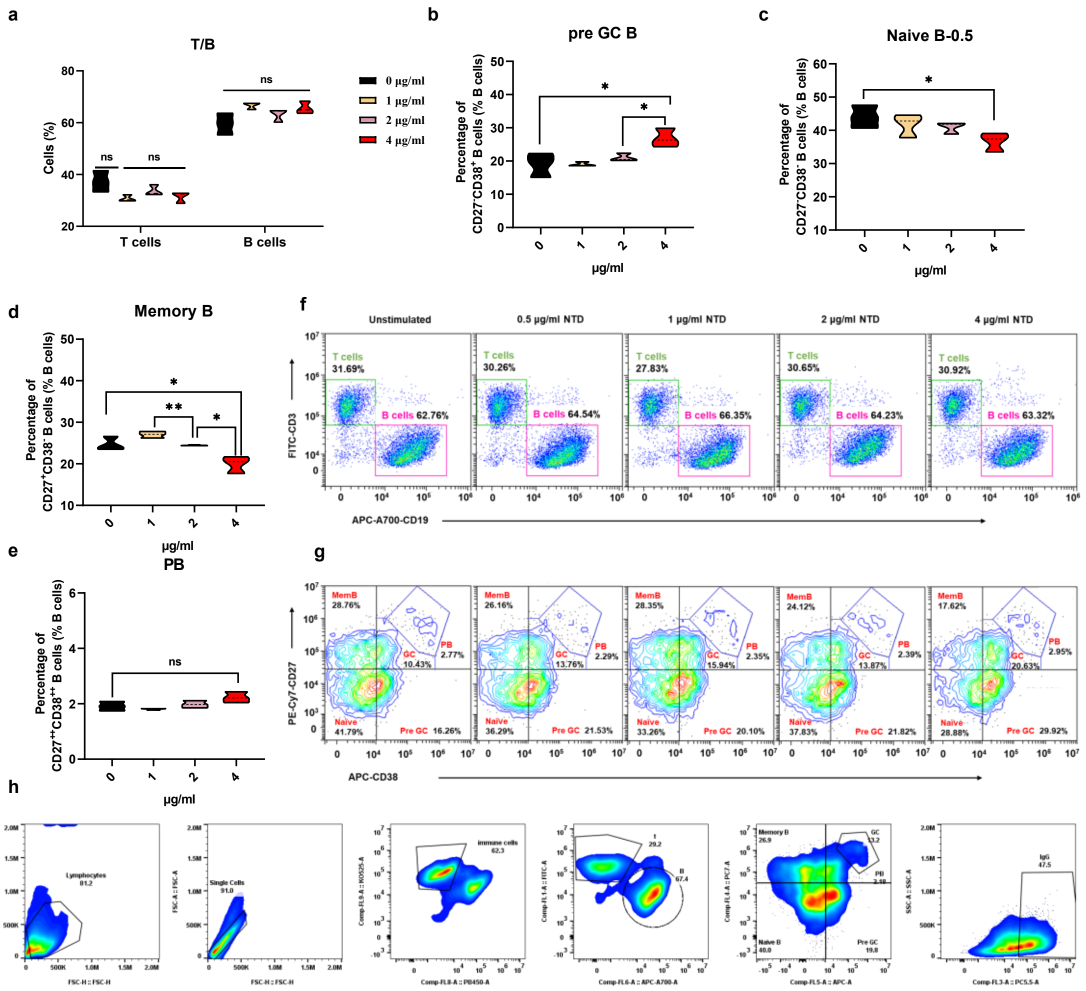

**Supplementary Fig. S3 | Flow cytometric characterization of antigen dose-dependent responses in tonsil immune organoids. a,** Frequency of total T cells and B cells across the indicated antigen doses after 7 days of culture. **b,** Frequency of pre-GC B cells across the indicated antigen doses. **c,** Frequency of naive B cells across the indicated antigen doses. **d,** Frequency of memory B cells across the indicated antigen doses. **e,** Frequency of plasmablasts across the indicated antigen doses. **f,** Representative flow cytometry gating plots showing total T-cell and B-cell populations across the indicated antigen doses. **g,** Representative quadrant gating plots used to define GC-associated cell populations. **h,** Sequential gating strategy used to identify germinal center–related B-cell populations in tonsil immune organoids after 7 days culture.

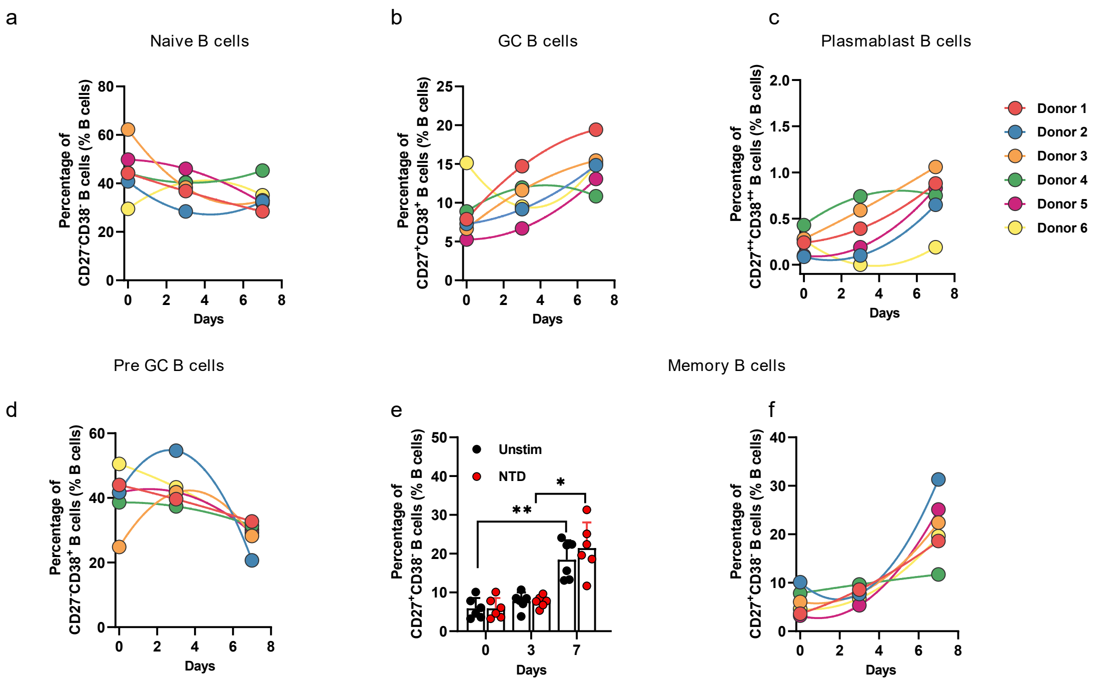


**Supplementary Fig. 4 | Additional characterization of temporal B cell remodeling in tonsil immune organoids. a,** Quantification of Naive B cells across days 0, 3, and 7 under unstimulated and antigen-stimulated conditions. **b–e,** Donor-level trajectories of GC B cells (**b**), plasmablasts (**c**), pre-GC B cells (**d**), and memory B cells (**e**) during culture. Data are shown as mean ± SEM with individual donors overlaid. Statistical analyses were performed using two-way repeated-measures ANOVA. GP: *P < 0.0332, **P < 0.0021, ***P < 0.0002; ns P > 0.1234, not significant.

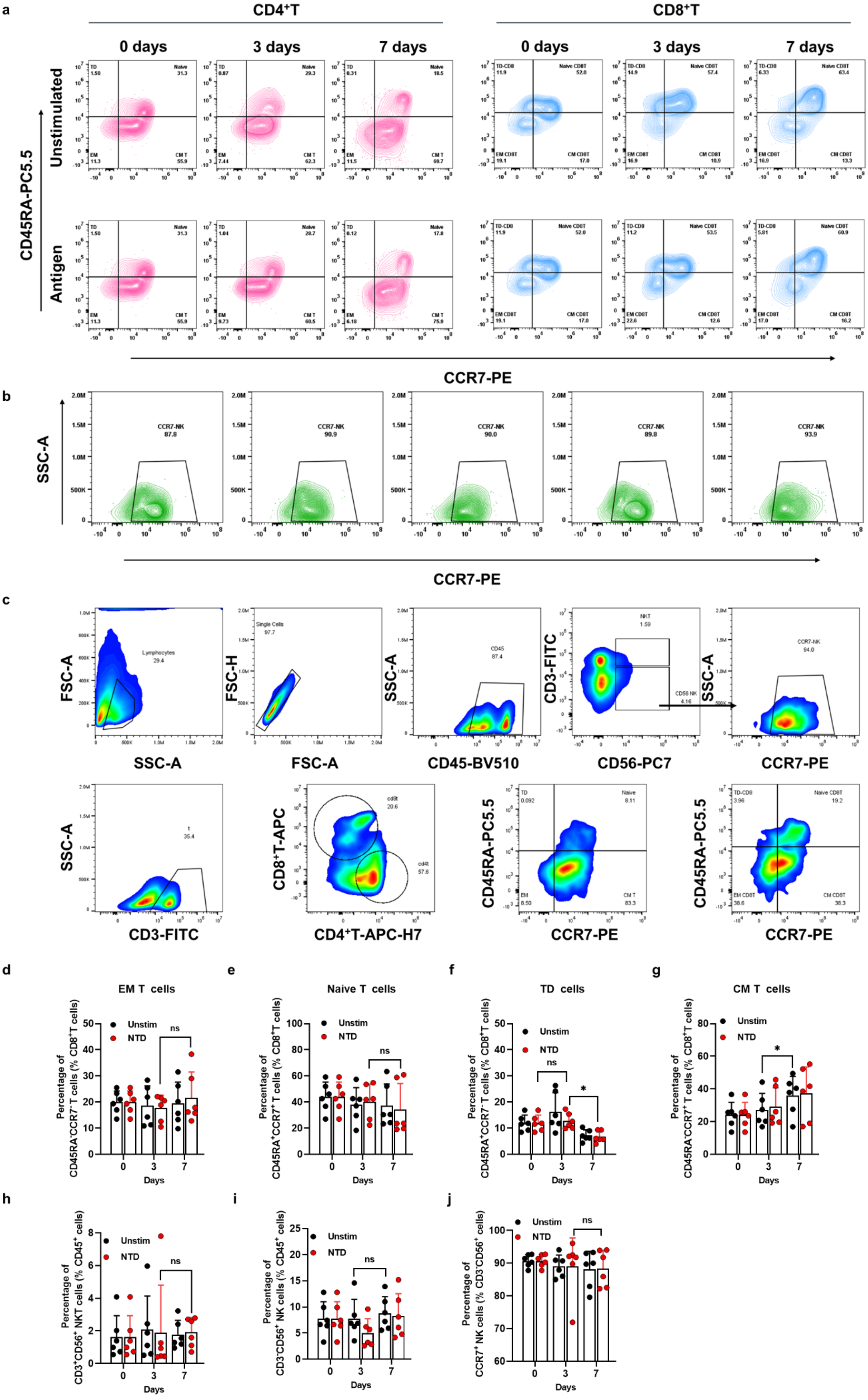
a
CD4+T
CD8+T
0 days
3 days
7 days
Unstimulated
Antigen
CD45RA-PC5.5
CCR7-PE
b
SSC-A
CCR7-NK
CCR7-PE
c
FSC-A
SSC-A
Lymphocytes
FSC-H
FSC-A
Single Cells
CD45-BV510
CD3-FITC
CD56-PC7
CCR7-PE
CD3-FITC
CD8+T-APC
CD4+T-APC-H7
CD45RA-PC5.5
d
EM T cells
e
Naive T cells
f
TD cells
g
CM T cells
Unstim
NTD
ns
Days
CD45RA-CCR7- T cells (% CD8+T cells)
CD45RA+CCR7+ T cells (% CD8+T cells)
CD45RA+CCR7- T cells (% CD8+T cells)
CD45RA-CCR7+ T cells (% CD8+T cells)
h
i
j
CD3+CD56+ NKT cells (% CD45+ cells)
CD3-CD56+ NK cells (% CD45+ cells)
CCR7+ NK cells (% CD3-CD56+ cells)

***Supplementary Fig. 5* | *A*dditional flow cytometric characterization of T-cell, DC-like, and NK-like remodeling in tonsil immune organoids. a,** Representative flow cytometry plots showing $CD4^{+}$T and $CD8^{+}$T cell differentiation states at days 0, 3, and 7 under unstimulated and antigen-stimulated conditions. **b,** Representative flow cytometry plots *showing frequency of* $CCR7^{+}$ *ce*lls within the $CD3^{-}CD56^{+}$ compartment over time. **c,** Sequential gating strategy used to identify $CD4^{+}$T, $CD8^{+}$T cell and NK like cells subsets in tonsil immune organoids. **d–g,** Quantification of $CD8^{+}$T cell subsets over time, including effector memory (EM) T cells (**d**), terminally differentiated (TD) T cells (**e**), central memory (CM) T cells (**f**), and naive T cells (**g**). **h,** Frequency of $CD3^{+}CD56^{+}$ cells among $CD45^{+}$ cells at days 0, 3, and 7 in unstimulated and NTD-stimulated organoids. j, Frequency of $CD3^{-}CD56^{+}$ cells among $CD45^{+}$ cells over time. i, Frequency of $CCR7^{+}$ cells within the $CD3^{-}CD56^{+}$ compartment over time. Data are shown as mean ± SEM with individual donors overlaid, unless otherwise indicated. Statistical analyses were performed using two-way repeated-measures ANOVA followed by Tukey's multiple comparisons test. ns, not significant; $*P < 0.05$, $**P < 0.01$, $***P < 0.001$, $****P < 0.0001$.

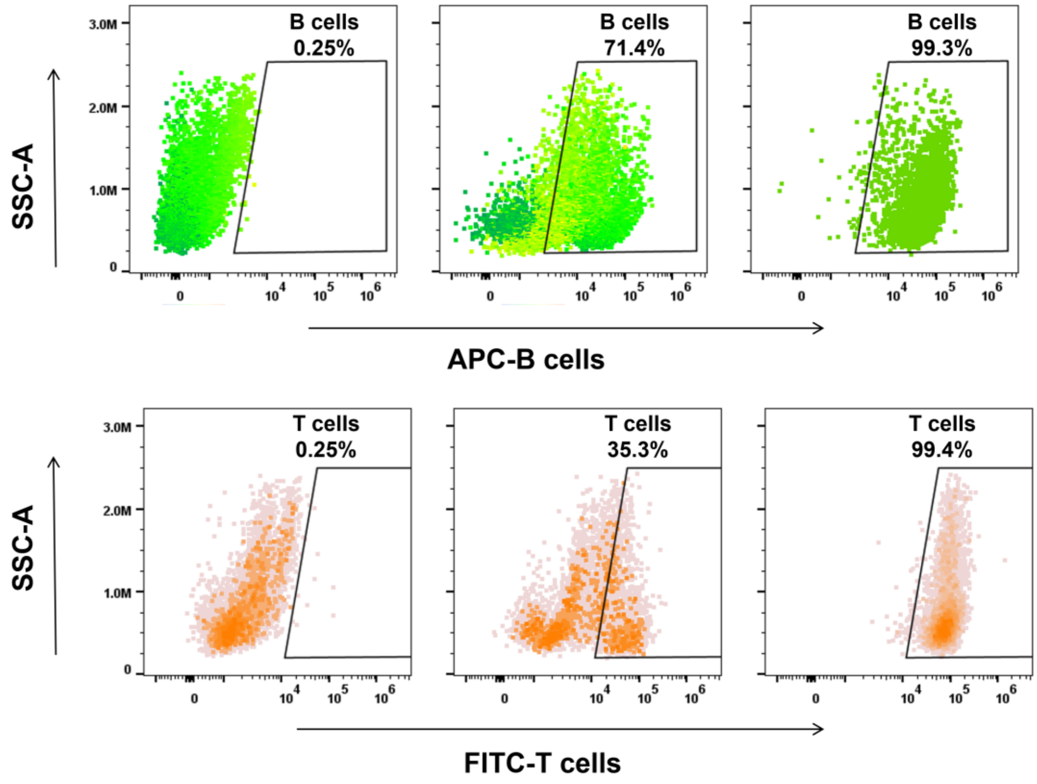


**Supplementary Fig. 6 |** T cells and B cells were isolated by magnetic separation prior to fluorescent labeling for live-imaging experiments. Representative flow-cytometric dot plots showing the enrichment of APC-B-positive B cells and FITC-T-positive T cells under the corresponding conditions. Numbers indicate the percentage of gated positive cells. These results verify the efficiency of the magnetic separation workflow used for subsequent time-lapse imaging of labeled lymphocytes in tonsil immune organoids.

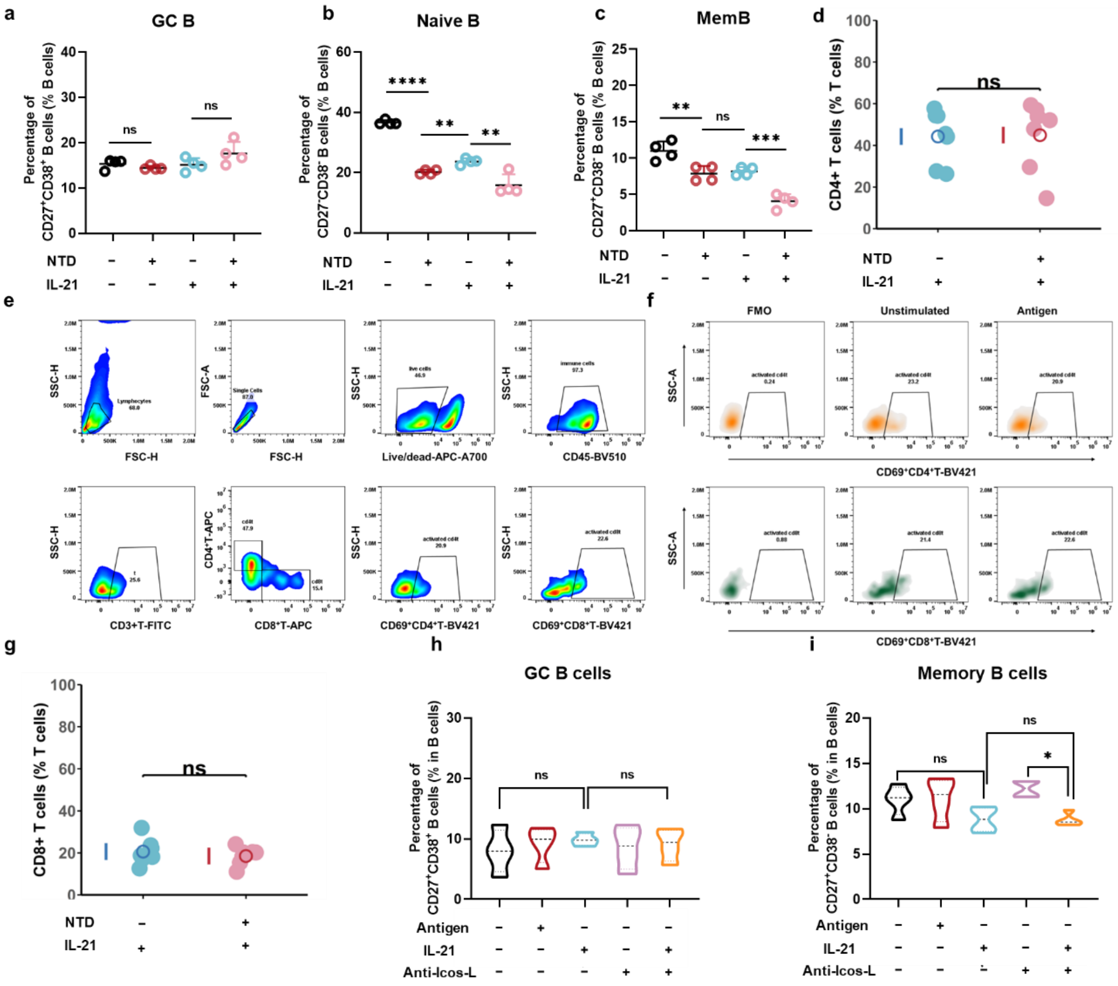


**Supplementary Fig. 7 | Additional flow-cytometric characterization of IL-21-associated responses and restricted ICOS-L signaling in tonsil immune organoids. a–c**, Quantification of GC B cells (**a**), naive B cells (**b**), and memory B cells (**c**) under the indicated conditions (unstimulated, NTD, IL-21, and NTD+IL-21) after 7 days of culture. **d–e**, Quantification of total CD4$^+$ T cells (**d**) and total CD8$^+$ T cells (**e**) in IL-21-containing cultures with or without NTD stimulation at day 7. **f–g,** Quantification of GC B cells (**f**) and memory B cells (**g**) across the indicated five conditions (unstimulated, NTD, IL-21, anti-ICOS-L, and IL-21+anti-ICOS-L) after 7 days of organoid culture. Data are shown as individual donors, with mean values indicated. Statistical comparisons were performed using unpaired two-tailed t-tests for the indicated comparisons. ns, not significant; *P < 0.05, **P < 0.01, ***P < 0.001, ****P < 0.0001.

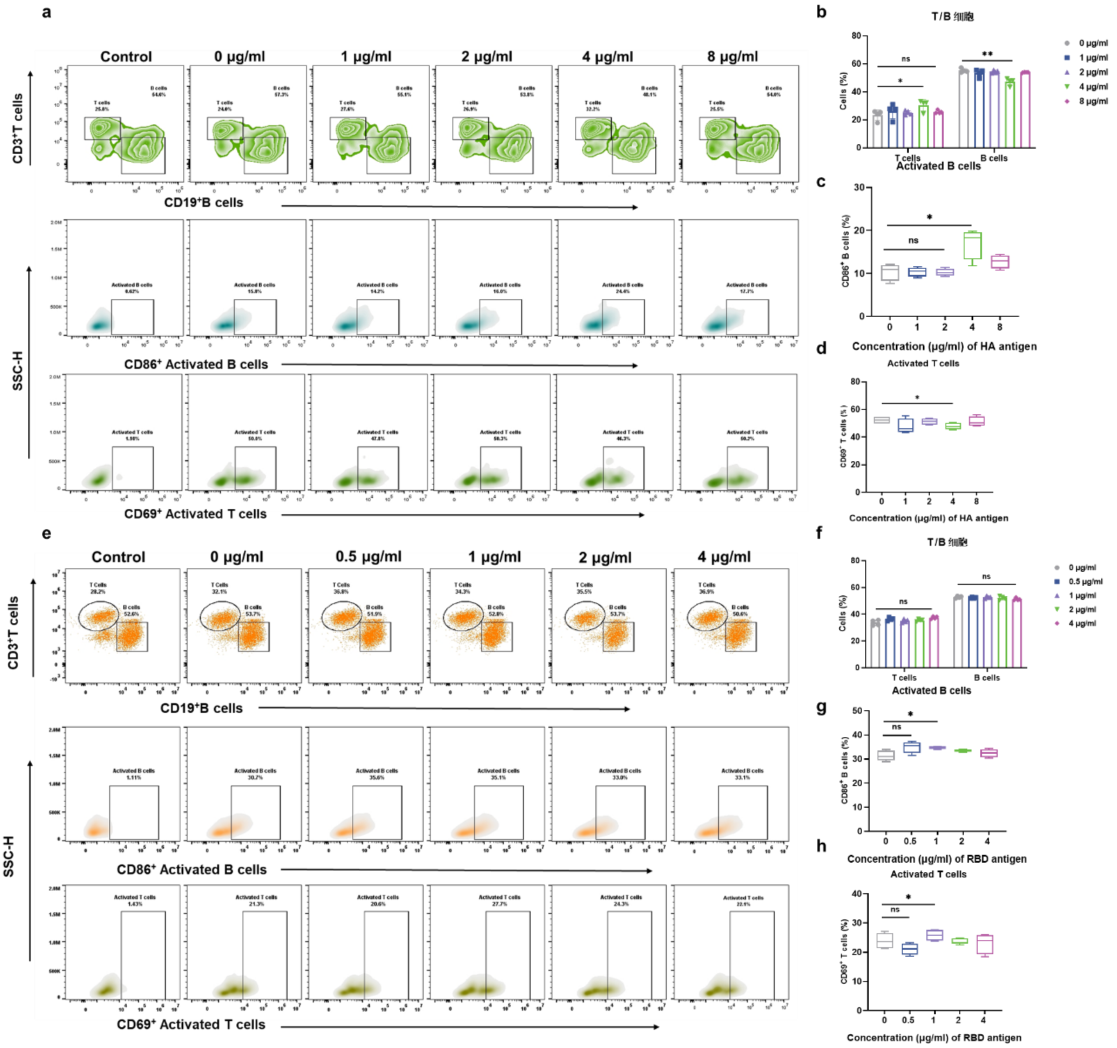


**Supplementary Fig. 8 | Optimization of antigen concentrations for short-term stimulation in tonsil immune organoids. a,** Representative flow-cytometry plots showing T-cell and B-cell distributions, together with gated CD86$^+$ activated B cells and CD69$^+$ activated T cells, after 24 h stimulation with increasing concentrations of HA antigen. **b,** Quantification of total T-cell and B-cell frequencies under the indicated HA concentrations. **c,** Quantification of CD86$^+$ activated B cells following 24 h HA stimulation. **d,** Quantification of CD69$^+$ activated T cells following 24 h HA stimulation. **e,** Representative flow-cytometry plots showing T-cell and B-cell distributions, together with gated CD86$^+$ activated B cells and CD69$^+$ activated T cells, after 24 h stimulation with increasing concentrations of RBD antigen. **f,** Quantification of total T-cell and B-cell frequencies under the indicated RBD concentrations. **g,** Quantification of CD86$^+$ activated B cells following 24 h RBD stimulation. **h,** Quantification of CD69$^+$ activated T cells following 24 h RBD stimulation. Together, these data were used to define organoid-compatible working concentrations for HA and RBD for subsequent vaccine-formulation studies. Data are shown as mean ± SEM with individual donors overlaid. Statistical significance was assessed using two-tailed paired or unpaired Student's t-tests, as specified in the corresponding panels.

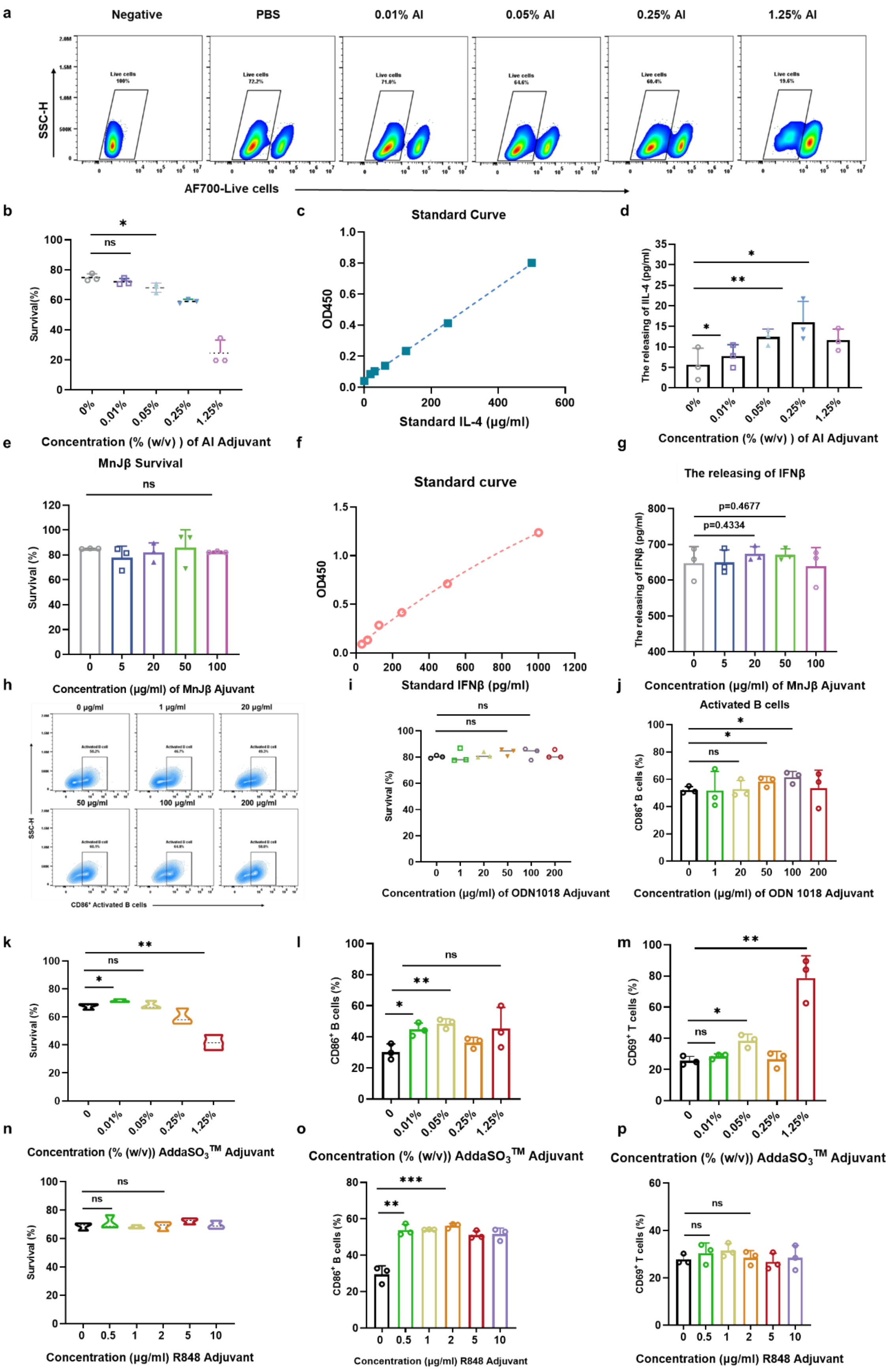


**Supplementary Fig. 9 | Optimization of working concentrations for clinically** tested

**adjuvants in tonsil immune organoids. a,** Representative viability gating plots following 24 h stimulation with increasing concentrations of Al. **b,** Quantification of organoid survival after stimulation with increasing concentrations of Al. **c,** Standard curve used for IL-4 quantification. **d,** Quantification of IL-4 release under increasing concentrations of Al. **e,** Quantification of organoid survival after stimulation with increasing concentrations of MnJβ. **f,** Standard curve used for IFNβ quantification. **g,** Quantification of IFNβ release under increasing concentrations of MnJβ. **h,** Representative flow-cytometry plots showing $CD86^+$ activated B cells after 24 h stimulation with increasing concentrations of ODN1018. **i,** Quantification of organoid survival after stimulation with increasing concentrations of ODN1018. **j,** Quantification of $CD86^+$ activated B cells under increasing concentrations of ODN1018. **k,** Quantification of organoid survival after stimulation with increasing concentrations of AddaSO3. **l.** Quantification of $CD86^+$ activated B cells under increasing concentrations of AddaSO3. **m,** Quantification of $CD69^+$ activated T cells under increasing concentrations of AddaSO3. **n,** Quantification of organoid survival after stimulation with increasing concentrations of R848. **o,** Quantification of $CD86^+$ activated B cells under increasing concentrations of R848. **p,** Quantification of $CD69^+$ activated T cells under increasing concentrations of R848. Across all adjuvants, working concentrations were selected by jointly considering preservation of organoid viability and induction of early immune activation. These experiments established organoid-compatible concentrations for downstream adjuvant and vaccine-formulation studies. Data are shown as mean ± SEM with individual donors overlaid. Statistical significance was assessed using two-tailed paired or unpaired Student's t-tests, as specified in the corresponding panels.

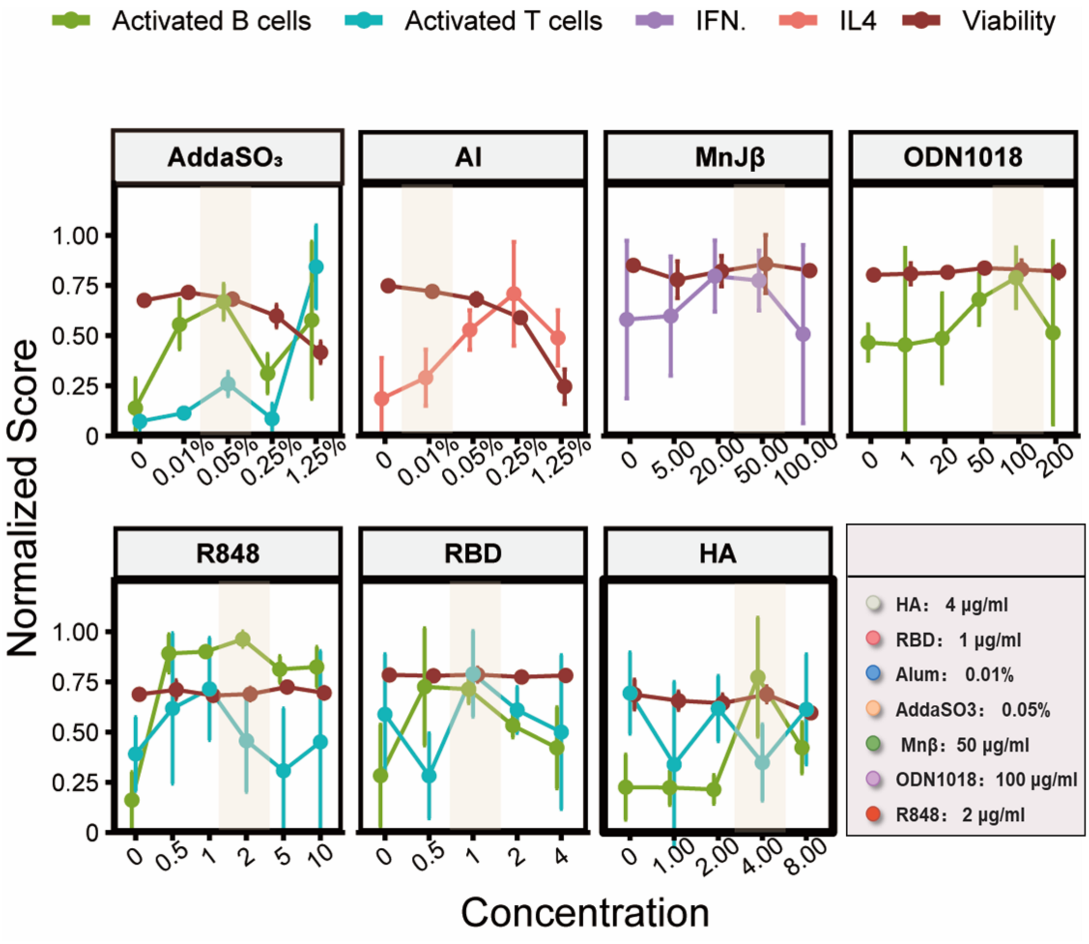

**Supplementary Fig. 10 | Optimization of working concentrations for clinically tested adjuvants in tonsil immune organoids.**

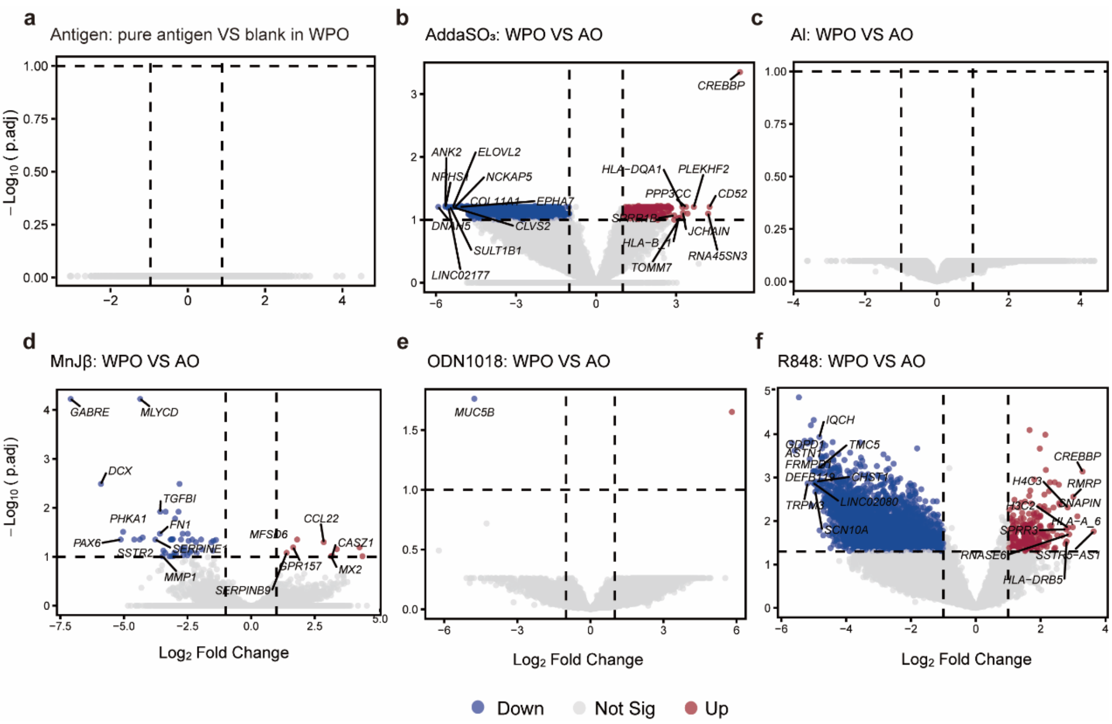


**Supplementary Fig. 11 | Volcano plots for adjuvant's different effect in WPOs and AOs. a,** Volcano plot showing the transcriptional effect of antigen stimulation alone in WPOs, generated by comparing the WPO no-adjuvant group (antigen only) with the blank group. Given the similar culture procedures used for WPOs and AOs, this comparison provides a reference for the antigen-only response in both models. **b-f,** Volcano plots comparing each adjuvant-treated condition between WPOs and Aos, illustrating model-dependent transcriptional differences in response to antigen plus adjuvant stimulation. For **a-f,** p values were adjusted using BH method. DEG cutoff were set as p.adj < 0.1 and $\log_2$Fold Change > 1.

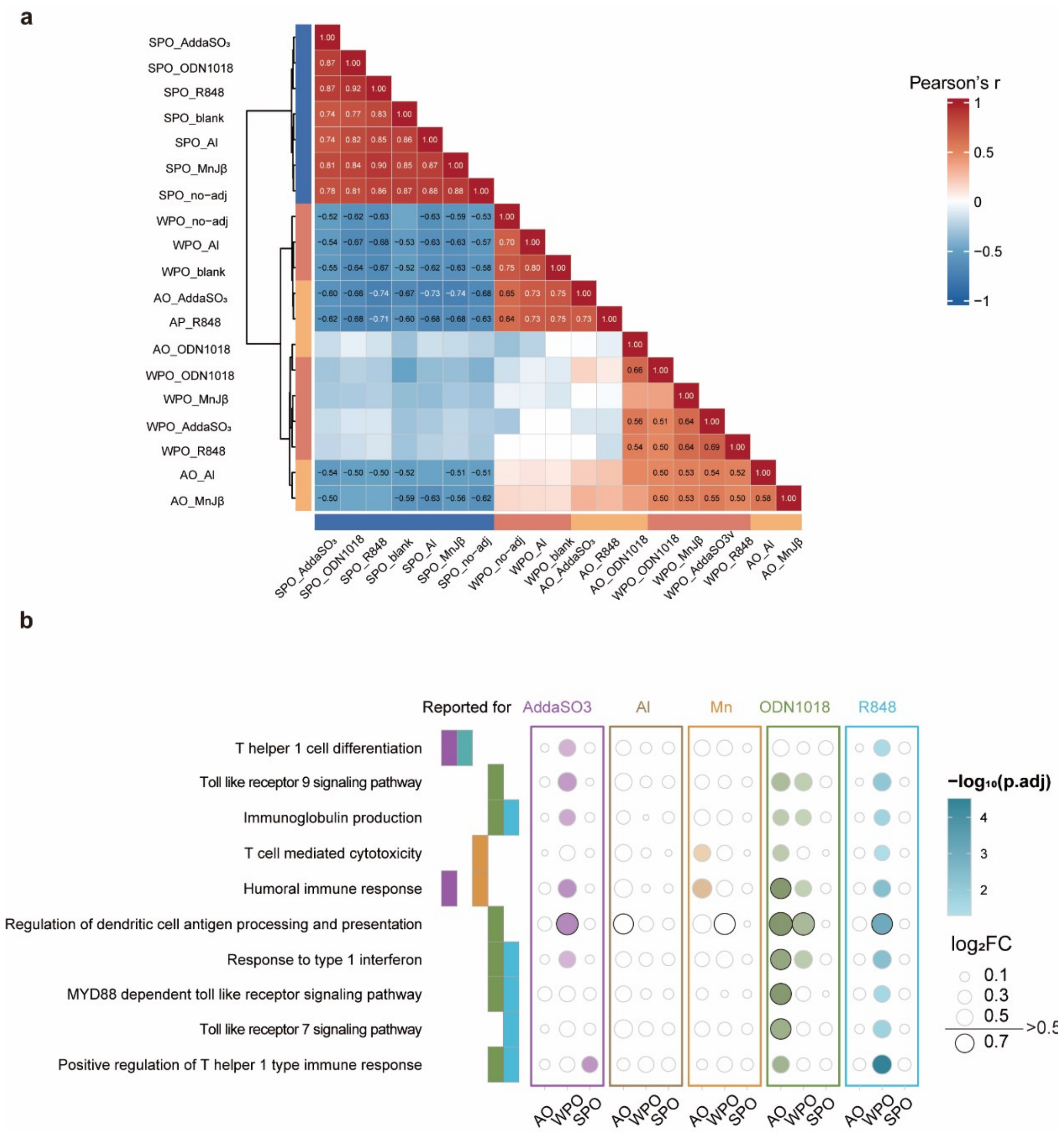


**Supplementary Fig. 12 | GSVA analysis of adjuvant-stimulated organoids. a,** Correlation heatmap based on GSVA enrichment scores. Only significant correlations ($p$.adj <0.05, | r |>0.5) are labeled, with color intensity representing Pearson's $r$. P values were adjusted using BH method. **b,** Reported pathways activation level for different adjuvants among different cohorts.

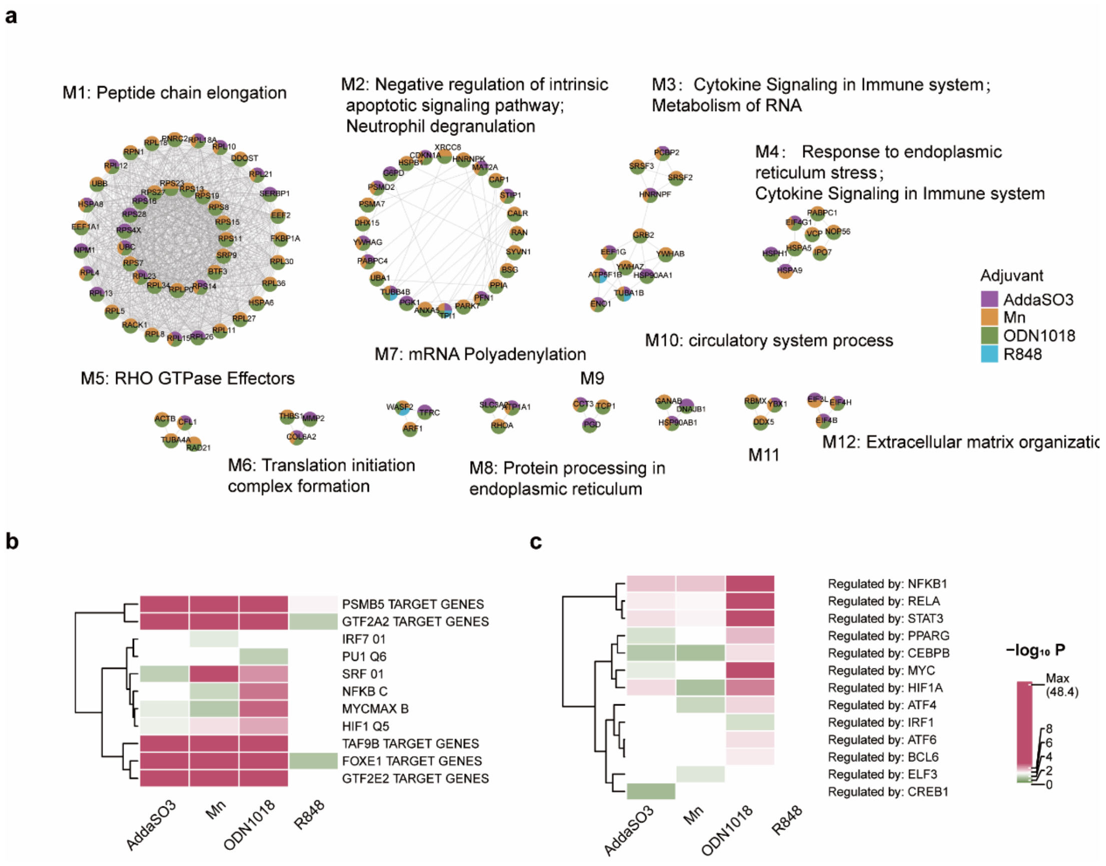


**Supplementary Fig. 13 | Conserved adjuvant-response genes revealed divergent and shared immune activation characteristics. a,** Core PPI networks identified using MCODE; **b-c,** TF analysis for regulator of conserved genes. In b, we used MsigDB C3: transcription factor targets as reference, while we used TRRUST in c. Both analysis achieved using Metascape.

**Supplementary Video 1 |** Time-lapse imaging of CellTracker 488-labeled T cells and DiD-labeled B cells in tonsil immune organoids under IL-21 treatment. Images were acquired every 10 min during the 24–48 h observation windows.

#### Key resources table

| REAGENT or RESOURCE | SOURCE | IDENTIFIER |
|---|---|---|
| Antibodies | | |
| BV510 Mouse Anti-Human CD45 (FC:1/50) | BD Biosciences | Cat# 563204 |
| FITC anti-human CD3 (FC:1/50) | BioLegend | Cat# 300440 |
| Alexa Fluor® 700 anti-human CD19 Antibody (FC:1/50) | BD Biosciences | Cat# 363034 |
| PE anti-human CD19 (FC:1/50) | BioLegend | Cat# 302208 |
| APC anti-human CD19 (FC:1/50) | BioLegend | Cat# 302212 |
| APC-H7 Mouse Anti-Human CD4 (FC:1/50) | BD Biosciences | Cat# 560158 |
| APC anti-human CD4 (FC:1/50) | BioLegend | Cat# 317318 |
| APC anti-human CD8 (FC:1/50) | | Cat# 344722 |

| | | |
|---|---|---|
| PE/Cyanine7 anti-human CD8 (FC:1/50) | BioLegend | Cat# 344750 |
| PE anti-human CD297 (PD-1) (FC:1/50) | BioLegend | Cat# 329905 |
| BV421 anti-human CD185 (CXCR5) (FC:1/50) | BioLegend | Cat# 356919 |
| APC Anti-human CD14 (FC:1/50) | BioLegend | Cat# 367118 |
| Anti-human IgG PC5.5 (FC:1/50) | | |
| BD Horizon™ BV510 Mouse Anti-Human CD27 (FC:1/50) | BD Biosciences | Cat# 563090 |
| PE/Cyanine7 anti-human CD27 (FC:1/50) | BioLegend | Cat# 356412 |
| APC anti-human CD38 Antibody (FC:1/50) | BioLegend | Cat# 303510 |
| BV421 Mouse Anti-Human CD86 (FC:1/50) | BD Biosciences | Cat# 562432 |
| BV421 CD69 (FC:1/50) | BioLegend | Cat# 310930 |
| PE anti-human CD69 (FC:1/50) | BioLegend | Cat# 310906 |
| PE anti-human CD197(CCR7) (FC:1/50) | BioLegend | Cat# 353204 |
| PerCP/Cyanine5.5 anti-human CD45RA (FC:1/50) | BioLegend | Cat# 304122 |
| PerCP-Cy 5.5 Mouse Anti-Human HLA-DR (FC:1/50) | BD Biosciences | Cat# 552764 |
| PE/Cyanine7 anti-human CD56(NCAM) (FC:1/50) | BioLegend | Cat# 362510 |
| Pan-Keratin Polyclonal antibody (FC:1/50) | proteintech | Cat# 26411-1-AP |
| Violet Fluorescent reactive dye 405 (FC:1/500) | Invitrogen | Cat# 2581659 |
| Horizon™ Fixable Viability Stain 700 (FC:1/500) | BD | Cat# 564997 |
| DAPI | MCE | Cat# HY-DY1081 |
| Anti-human CD20 Monoclonal Antibody (L26), eFluor™ 660 (IF:1/200) | eBioscience™ | Cat# 50-0202-82 |
| Purified anti-human CXCR4 (CD184) (IF:1/500) | BioLegend | Cat# 306501 |
| Human CD83 recombinant rabbit antibody (IF:1/500) | abcam | Cat# ab205343 |
| Recombine ant Anti-CD3 antibody (Rabbit mAb) (IF:1/200) | Servicebio | GB150004 |
| Recombine ant Anti-CD19 antibody (Mouse mAb) (IF:1/200) | Servicebio | GB15061 |
| F(ab')2-Goat anti rabbit IgG (H+L) Alexa FluorTM plus 488 (IF:1/1000) | Invitrogen | Cat# A48282TR |
| F(ab')2-Goat anti mouse IgG (H+L) Alexa Fluor™ plus 555 (IF:1/1000) | Invitrogen | Cat# A48287 |
| Prezalumab (Anti-cosI-L) | Sparkjade | Cat# SJ-BA1137 |
| Bacterial and virus strains | | |
| n/a | n/a | Cat# n/a |
| Biological samples | | |

| | | |
|---|---|---|
| Adult tonsil tissue | Peking University Shenzhen Hospital; https://www.pkuszh.com/ENGLISH | Cat# n/a |
| Chemicals, peptides, and recombinant proteins | | |
| Tissue storage solution | Miltenyi | Cat# 130-100-008 |
| RPMI Medium 1640 (1X) | Gibco™ | Cat# A10491 |
| DMEM (High glucose) | Cytiva | Cat# SH30022.01 |
| Fetal bovine serum (FBS) | Gibco™ | Cat# 10270-106 |
| Penicillin-Streptomycin (PS) | Gibco™ | Cat# 15140122 |
| Phosphate-buffered saline PBS (1×) | Gibco™ | Cat# 20-012-027 |
| Trypsin-EDTA (0.25%), phenol red | Gibco™ | Cat# 25200072 |
| Insulin-Transferrin-Selenium-Pyruvate (ITS-A) (100×) | Gibco™ | Cat# 51300044 |
| MEM (Non-essential Amino acid) | Gibco™ | Cat# 11140050 |
| GlutaMAX™ | Gibco™ | Cat# 35050061 |
| HEPES (1 M) | Gibco™ | Cat# 15630-080 |
| Y-27632 (ROCK inhibitor) | MCE | Cat# HY-10071 |
| Normocin™ | InvivoGen | Cat# ant-nr-05 |
| Nicotinamide | InvivoGen | Cat# HY-B0150 |
| L-Ascorbic acid (Vc) | Solarbio | Cat# 50-81-7 |
| A83-01 | MCE | Cat# HY-10432 |
| Human recombinant IL-21 | PeproTech | Cat# 200-21-10UG |
| Recombinant human BAFF | GenScript | Cat# Z02976-1 |
| Collagenase Type I, Cls I | Sigma-Aldrich | Cat# C1-28-100MG |
| Collagenase Type II, Cls I | Sigma-Aldrich | Cat# C2-28-100MG |
| DNase I | Beyotime | Cat# D7076 |
| DMSO | Sigma-Aldrich | Cat# C6295 |
| Red blood cell lysis buffer | Solarbio | Cat# R1011 |
| Matrigel Matrix (Growth Factor Reduced) | Corning | Cat# 356234 |
| Trypan Blue (0.4%) | Sigma-Aldrich | Cat# 72-57-1 |
| Iodophor | LIRCON | Cat# XY0058 |
| G418 | ThermoFisher | Cat# 10131035 |
| Hygromycin B | ThermoFisher | Cat# 10687010 |
| SuperKine™ Enhanced Antifade Mounting Medium with DAPI | Abbkine | Cat# BMU107 |
| Goat serum | Abbkine | Cat# BMS0050 |
| Sucrose | Solarbio | Cat# 57-50-1 |
| SARS-CoV-2 Spike N-Terminal Domain (NTD) | Sino Biological | Cat# 40591-V49H |
| Influenza A H3N2 (A/Darwin/9/2021) HA protein | Sino Biological | Cat# 40859-V08B |

| | | |
|---|---|---|
| SARS-CoV-2 (BA.2.12.1) Spike RBD protein | Sino Biological | Cat# 40592-V08H132-B |
| FcX | Biolegend | Cat# 422302 |
| CMFDA CellTracker | Yeasen | Cat# 40721ES50 |
| Biotinylated anti-CD3 antibody | Biolegend | Cat# 300304 |
| Streptavidin MicroBeads | Miltenyi | Cat# 130-048-102 |
| Critical commercial assays | | |
| AM/PI Live/Dead Viability Assay Kit | Beyotime | Cat# C2015M |
| EasySep Human B Cell Isolation Kit | Stemcell™ | Cat# 50102 |
| Cell membrane far-infrared fluorescence staining kit (DiD) | Beyotime | Cat# C1995S |
| Experimental models: Cell lines | | |
| L-WRN cells (secreting Wnt3a, R-spondin 3, Noggin) | ATCC | Cat# CRL-2647 |
| Deposited data | | |
| Bulk RNA Sequence (Raw Data) | | Accession |
| Software and algorithms | | |
| FlowJo (v10.8.1) | BD Biosciences | RRID:SCR_008520 |
| GraphPad Prism (v10) | GraphPad | RRID:SCR_002798 |
| ImageJ (v1.54f) | NIH | RRID:SCR_003070 |
| R (4.3.1) | R-Project | RRID:SCR_001905 |
| ggplot2 (v3.4.3) | R Package | RRID:SCR_014601 |
| BioRender | Biorender | RRID:SCR_018361 |
| Other | | |
| 70 µm Cell Strainer | Falcon | Cat# 352350 |
| Pasteur pipette | JETBIOFIL | Cat# PP102030 |
| 60 cm² TC-treated Culture Dish | Corning | Cat# 430166 |
| 100 cm² TC-treated Culture Dish | Corning | Cat# 430167 |
| BeyoCool™ Cell Freezing Container | Beyotime | Cat# FCFC012 |
| Metal Ice Box | Biosharp | Cat# BC032 |
| Cryogenic vials (2 mL) | NEST | Cat# 607401 |
| 50 mL Centrifuge tube | Corning | Cat# 430790 |
| 15 mL Centrifuge tube | Corning | Cat# 430828 |
| 5 mL syringe plunger | Gufastore | Cat# GF22101402 |
| 0.4 µm transwell PET | Corning | Cat# 3470 |
| Laser confocal dish | Biosharp | Cat# BS-15-GJM |
| Neg-50 Frozen Section Medium（OCT） | ThermoFisher | Cat# 6502B |
| $CO_2$ Incubator (37℃, 5% $CO_2$) | ESCO | Model# CCL-050B-8 |

| | | |
|---|---|---|
| Water bath | Thermo Scientific | Cat# TSGP02 |
| Confocal laser scanning microscope | Olympus Corporation | Model# FV3000 |
| CytoFLEX Flow Cytometer | Beckman | Model# V0-B5-R3 |

#### Supplementary Table 1 for tissue donor characteristics

| Donor | Age (years) | Sex | Indication for surgery | COVID-19 vaccination | COVID-19 Infection history |
|---|---|---|---|---|---|
| 1 | 47 | F | hypertrophy | 2 doses | Yes |
| 2 | 36 | M | hypertrophy | 2 doses | Yes |
| 3 | 28 | F | hypertrophy | 2 doses | Yes |
| 4 | 36 | M | hypertrophy | 2 doses | Yes |
| 5 | 27 | M | hypertrophy | 2 doses | Yes |
| 6 | 46 | M | hypertrophy | 3 doses | Yes |
| 7 | 18 | M | Recurrent tonsillitis | 3 doses | Yes |
| 8 | 29 | M | hypertrophy | 2 doses | Yes |
| 9 | 29 | M | hypertrophy | 3 doses | Yes |
| 10 | 31 | F | Recurrent tonsillitis | 3 doses | Yes |
| 11 | 38 | M | hypertrophy | 3 doses | No |
| 12 | 28 | M | Recurrent tonsillitis | 2 doses | Yes |
| 13 | 25 | M | Recurrent tonsillitis | 3 doses | Yes |
| 14 | 34 | F | Recurrent tonsillitis | 2 doses | Yes |
| 15 | 19 | M | hypertrophy | 2 doses | Yes |
| 16 | 36 | M | Recurrent tonsillitis | 2 doses | Yes |
| 17 | 40 | M | hypertrophy | 3 doses | Yes |
| 18 | 18 | M | hypertrophy | 3 doses | Yes |
| 19 | 29 | M | hypertrophy | 2 doses | Yes |
| 20 | 25 | M | hypertrophy | 3 doses | Yes |
| 21 | 36 | F | hypertrophy | 3 doses | Yes |
| 22 | 32 | M | hypertrophy | 3 doses | Yes |
| 23 | 28 | F | hypertrophy | 3 doses | Yes |

Table note: All specimens were derived from palatine tonsils obtained at tonsillectomy. Original cryostorage identifiers were replaced with anonymized sample IDs.

**Supplementary Table 2 Flow antibody panels**

| Marker | Fluorochrome | Dilution |
|---|---|---|
| B-cell differentiation and class-switching panel | | |
| Violet Fluorescent reactive dye 405 | 405 | 1:500 |
| CD45 | BV510 | 1:50 |
| CD19 | APC-A700 | 1:50 |
| CD3 | FITC | 1:50 |
| CD38 | APC | 1:50 |
| CD27 | PC7 | 1:50 |
| IgG | PC5.5 | 1:50 |
| FcX | - | 1:50 |
| T-cell differentiation and lymphoid homing panel | | |
| CD45 | BV510 | 1:50 |
| CD3 | FITC | 1:50 |
| CD4 | APC-H7 | 1:50 |
| CD8 | APC | 1:50 |
| CD56 | PC7 | 1:50 |
| CD45RA | PC5.5 | 1:50 |
| CCR7 | PE | 1:50 |
| FcX | - | 1:50 |
| DC-like and B-cell maturation staining panel | | |
| CD45 | BV510 | 1:50 |
| CD3 | FITC | 1:50 |
| CD19 | APC-A700 | 1:50 |
| CD14 | APC | 1:50 |
| CD56 | PC7 | 1:50 |
| HLA-DR | PC5.5 | 1:50 |
| DAPI | DAPI | 1:1000 |
| FcX | - | 1:50 |
| T-cell activation staining panel | | |
| Horizon™ Fixable Viability Stain 700 | APC-A700 | 1:500 |
| CD45 | BV510 | 1:50 |
| CD3 | FITC | 1:50 |
| CD4 | APC | 1:50 |
| CD8 | PC7 | 1:50 |
| CD69 | BV421 | 1:50 |

| | | |
|---|---|---|
| FcX | - | 1:50 |
| B-cell activation staining panel | | |
| Horizon™ Fixable Viability Stain 700 | | 1:500 |
| CD45 | BV510 | 1:50 |
| CD19 | PE | 1:50 |
| CD3 | FITC | 1:50 |
| CD86 | BV421 | 1:50 |
| IgG | PC5.5 | 1:50 |
| FcX | - | 1:50 |
| B-cell activation and differentiation panel | | |
| CD3 | FITC | 1:50 |
| CD19 | APC-A700 | 1:50 |
| CD86 | BV421 | 1:50 |
| CD38 | APC | 1:50 |
| CD27 | BV510 | 1:50 |
| PI | PC5.5 | 1:1000 |
| FcX | - | 1:50 |
| Tfh-like cell staining panel | | |
| Horizon™ Fixable Viability Stain 700 | APC-A700 | 1:500 |
| CD45 | BV510 | 1:50 |
| CD3 | FITC | 1:50 |
| CD4 | APC | 1:50 |
| PD-1 | PE | 1:50 |
| CXCR5 | BA421 | 1:50 |
| FcX | - | 1:50 |
| T-cell, B-cell, and APC activation panel | | |
| Horizon™ Fixable Viability Stain 700 | APC-A700 | 1:500 |
| CD3 | FITC | 1:50 |
| CD19 | APC | 1:50 |
| HLA-DR | PC5.5 | 1:50 |
| CD86 | BV421 | 1:50 |
| CD69 | PE | 1:50 |
| FcX | - | 1:50 |

**Supplementary Table 3 Composition of complete tonsil organoid medium**

| Reagent | Final concentration |
|---|---|

| | |
|---|---|
| RPMI Medium 1640 (1X) | 59% (v/v) |
| L-WRN conditioned medium (Wnt3a, R-spondin 3, Noggin) | 25% (v/v) |
| FBS | 10% (v/v) |
| PS | 1% (v/v) |
| GlutaMAX™ | 1× |
| ITS-A (100×) | 1× |
| MEM non-essential amino acids (NEAA) | 1× |
| Y-27632 | 10 μM |
| HEPES | 10 mM |
| Nicotinamide | 10 mM |
| Vc | 20 μg/ml |
| A83-01 | 20 nM |
| Human recombinant IL-21 | 10 ng/mL |
| Recombinant human BAFF | 100 ng/mL |
| Normocin™ | 0.1mg/mL |
| NTD Antigen | 4 μg/mL |
| *(HA Antigen)* | 4 μg/mL |
| *(RBD Antigen)* | 1 μg/mL |

### Methods

#### *Human tonsil specimens*

All human palatine tonsils samples used in this study were collected with written informed consent under protocols approved by the Research Ethics Committee of Peking University Shenzhen Hospital, the General Hospital of Southern Theater Command of PLA (Guangzhou Liuhuaqiao Hospital) and the Ethics Review Committee of Tsinghua Shenzhen International Graduate School (**Ethics Approval Number: 2024F103**). All experiments were performed in accordance with institutional and national guidelines and regulations governing the use of human biological materials. In this study, the participant cohort consisted of tissues acquired from both male and female individuals aged 18-47 years (**Supplementary Table 1**).

#### *Tonsil tissue dissociation and preparation of single-cell suspensions*

Fresh tonsil specimens were placed immediately into sterile tissue storage solution (Miltenyi) after surgical removal and transported on ice to the laboratory for processing as soon as possible. Tonsil tissues were processed using a staged dissociation workflow adapted from published human tonsil organoid protocols[1, 2] and further optimized for downstream organoid culture in our system. Briefly, visibly necrotic tissue and non-lymphoid debris were removed under sterile conditions. Tissue fragments were then decontaminated by brief povidone–iodine rinsing for 20 s, followed by washing in sterile PBS (Gibco™). After decontamination, samples were subjected to a 20-min antimicrobial pre-treatment on a metal ice brick using antibacterial buffer containing 2% fetal bovine serum (FBS) (Gibco™), penicillin–streptomycin, Normocin (0.1 mg/mL) (InvivoGen), and Y-27632 (1 μM) (MCE). After pre-processing, tissues were minced into small fragments and transferred into digestion medium prepared in DMEM and supplemented with collagenase I (0.5 mg/mL) (Sigma-Aldrich), collagenase II (0.5 mg/mL) (Sigma-Aldrich), DNase I (50 μg/mL) (Beyotime), Y-27632 (1 μM), and Normocin (0.1 mg/mL). An initial round of enzymatic digestion was carried out at 37 °C with gentle agitation at 70 rpm for 20 min. Following the first digestion, immune cells were gently released by mechanical trituration, and the supernatant containing liberated cells was collected. Residual tissue fragments were then returned to fresh digestion medium and subjected to a second round of enzymatic digestion for an additional 30–60 min to maximize cell recovery. Supernatants from both digestion steps were pooled and passed through a 70-μm cell strainer. Filtered cell suspensions were washed in PBS-based buffer and treated with red blood cell lysis buffer when necessary. Cells were then pelleted, resuspended in PBS, counted, and assessed for viability by trypan blue exclusion before cryopreservation in DMSO-containing freezing medium. Unless otherwise specified, all subsequent handling steps were performed at low temperature.

#### *Generation and culture of tonsil immune organoids*

Where cryopreserved samples were used, vials were rapidly thawed in a 37 °C water bath, washed in PBS, and resuspended in organoid medium before cell counting and plating. Complete tonsil organoid medium was prepared as summarized in **Supplementary Table 3** and consisted of RPMI 1640 supplemented with L-WRN conditioned medium, fetal bovine serum, penicillin–streptomycin, GlutaMAX, ITS-A, MEM supplement, HEPES, Y-27632, nicotinamide, vitamin C, A83-01, recombinant human IL-21, recombinant human BAFF, and Normocin. L-WRN conditioned medium was prepared as previously described and used as a source of Wnt3a, R-spondin 3, and Noggin[3]. For organoid assembly, cells were resuspended in ice-cold complete tonsil organoid medium containing 5% (v/v) Matrigel and adjusted to a final density of 2 × 10^6 viable cells per 100 μL. Cell–matrix suspensions were gently mixed and seeded into 0.4-μm 24-pole transwell inserts, while 600 μL of complete organoid medium was added to the lower chamber. Before seeding, transwell plates were equilibrated at 37 °C in 5% $CO_2$. To preserve matrix integrity and cell viability, Matrigel-containing suspensions were kept on ice during preparation and seeded immediately after mixing. Organoids were maintained at 37 °C in a humidified 5% $CO_2$ incubator, and the medium in the lower chamber was replaced every 2 days unless otherwise specified. For antigen-priming experiments, antigen was added only once at the time of plating and was not replenished during subsequent medium changes. NTD model antigen was used at a final concentration of 4 μg/mL during initial seeding. Under these conditions, germinal center-like structures became detectable from approximately day 7 and continued to mature thereafter. For adjuvant evaluation experiments, organoids were first primed with HA (4 μg/mL) or RBD (1 μg/mL) antigen during plating and cultured for 7 days. On day 8, cultures were reassigned to no-antigen, antigen-only, or antigen-plus-adjuvant conditions using the predefined working concentrations described in the Results. For short-term experiments, organoids were harvested after 24 h of stimulation. For prolonged-phase experiments, cultures were maintained for an additional 7 days, with medium replacement every 2 days, before downstream analysis.

#### *Flow cytometry*

For flow-cytometric analysis, tonsil organoids were harvested and dissociated into single-cell suspensions. Briefly, medium from the upper chamber was collected, Cells were then gently dissociated by pipetting, combined with PBS washes from the insert, and passed through a 100-μm cell strainer to obtain single-cell suspensions. After centrifugation at 500 × g for 5 min, cell pellets were resuspended in staining buffer for downstream antibody labeling.
For viability assessment, cells were first stained with a protein-binding live/dead dye (1:500) for 20 min at 4 °C in the dark, followed by one PBS wash. Cells were then incubated with fluorochrome-conjugated antibodies for surface staining for 20 min at 4 °C in the dark following the addition of antibody cocktails (**Table S2**). All antibody solutions were diluted using Staining Buffer and Human TruStain FcX to prevent nonspecific antibody binding. Multiple multicolor panels were used in this study according to the experimental readout, including panels for immune composition, B-cell activation and germinal-center-associated phenotypes, B-cell and dendritic-cell maturation, T-cell activation, T-cell differentiation, and NK-like cell phenotyping. These panels included markers for $CD45^+$ leukocytes, $CD3^+$ T cells, $CD4^+$ and $CD8^+$ T-cell subsets, $CD19^+$ B cells, $CD56^+$ NK-like cells, and HLA-$DR^+$dendritic-cell-associated populations. B-cell-focused panels further included activation- and germinal-center-related markers such as CD86, HLA-DR, CXCR4, whereas T-cell-focused panels included activation and differentiation markers such as CD69, CCR7, and CD45RA. Additional panels were used to assess HLA-DR-associated maturation in B-cell and dendritic-cell compartments, as well as CD56-based phenotyping of NK-like populations.
Because individual panels were optimized for distinct readouts, some markers were paired with different fluorophores across panels. Full antibody information, including marker, fluorophore, vendor, catalog number, and dilution, is provided in **Key Resources Table and Supplementary Table 2**. Samples were acquired on a Beckman Coulter CytoFLEX flow cytometer and analyzed

using FlowJo v10. Debris, doublets, and dead cells were excluded before downstream gating. Flow cytometry data were analyzed using FlowJo.

##### *ELISA*

Culture supernatants were collected at the indicated time points and clarified by centrifugation at 3000 × g for 5 min. The supernatants were then carefully transferred, freeze-dried using a lyophilizer, and stored at −80 °C until analysis. Before ELISA, the lyophilized samples were reconstituted in 200–300 μL of the corresponding sample diluent. Levels of IL-4, IFNβ, and antigen-specific IgG against SARS-CoV-2 RBD were quantified using commercially available ELISA kits according to the manufacturers' instructions. Standards and samples were assayed in duplicate or triplicate. Absorbance was measured at 450 nm using a microplate reader, with a reference wavelength when applicable. Concentrations were calculated from standard curves generated by four-parameter logistic regression or linear regression, according to the kit instructions.

##### *Immunofluorescence staining and confocal microscopy*

For fixed-tissue imaging, organoids were collected and fixed in 2% paraformaldehyde overnight or 4 h at 4 °C. Samples were washed in PBS, cryoprotected in 10%, 20% and 30% sucrose, embedded in OCT, and sectioned at 10–20 μm using a cryostat. Sections were permeabilized in 0.3% Triton X-100, blocked in 5% goat serum for 2h on the room tempature. Then Sections were incubated with primary antibodies against markers of interest, including CD83, CXCR4, CD19 PD-1, CXCR5 and CD3, followed by species-appropriate fluorophore-conjugated secondary antibodies, including F(ab')2-Goat anti rabbit IgG (H+L) Alexa FluorTM plus 488 or F(ab')2-Goat anti mouse IgG (H+L) Alexa FluorTM plus 555, for 1 h at 20°C-25°C in the dark. Add Tertiary antibody (Anti-human CD20 Monoclonal Antibody (L26), eFluor$^{TM}$ 660) incubation solution to each section and incubate 3h at 20°C-25°C in a humidified chamber.

Nuclei were counterstained with DAPI. Images were acquired using Nikon A1 confocal microscope with identical acquisition settings within each experiment. Image analysis was performed in Fiji/ImageJ, and quantitative measurements such as area, mean fluorescence intensity, or compartment-specific signal were extracted using predefined analysis pipelines.

##### *Time-lapse imaging of T- and B-cell dynamics from organoid*

Tonsil-derived single-cell suspensions were used for magnetic isolation of T and B cells prior to live imaging. T cells were positively selected using biotinylated anti-CD3 antibody and Streptavidin MicroBeads, whereas B cells were isolated by negative selection using an EasySep Human B Cell Isolation Kit. Isolated T cells were labeled with CellTracker 488, and isolated B cells were labeled with DiD. Labeled T and B cells were then recombined with the remaining unlabeled tonsil cells and seeded into detachable 12-well Transwell inserts for organoid culture.

For dynamic imaging, Transwell inserts were transferred to glass-bottom dishes containing organoid medium in the chamber at 37 °C and 5% $CO_2$ and imaged on an inverted fluorescence microscope under NTD + IL-21 conditions. Images were acquired every 10 min over two time windows, 0–16 h and 24–48 h, using fluorescence channels for CellTracker 488 and DiD to visualize T-cell and B-cell movement, respectively. Time-lapse image series were compiled into videos for subsequent analysis. Laser power, exposure time, gain, and imaging interval were kept constant within each experiment to minimize photobleaching and phototoxicity.

#### *RNA-sequencing Data Processing and Quantification*

Raw sequencing data in FASTQ format were initially subjected to comprehensive quality control using FastQC (v0.12.1) to assess read quality and identify potential sequencing artifacts. To ensure high-quality input for downstream alignment, adapter sequences and low-quality bases were trimmed using cutadapt (v4.4)[4].

The resulting clean reads were subsequently mapped to the human reference genome *(GRCh38.p14)* using the highly efficient, splice-aware aligner STAR (v2.7.10b)[5]. To guide the spliced alignments and ensure accurate genomic feature identification, the corresponding NCBI RefSeq gene annotation file (GCF_000001405.40_GRCh38.p14_genomic.gtf) was utilized during the index building and mapping steps. Finally, the quantification of mapped reads was performed using featureCounts (v2.0.6)[6]. Read counts were aggregated at the gene level based on the RefSeq annotations, and only uniquely mapped reads were considered for the generation of the raw gene expression count matrix, which was subsequently used for downstream analyses.

To filter lowly expressed genes, raw counts were transformed using the rlog function in the DESeq2 R package[7]. Genes with a 90th percentile of rlog-transformed expression > 1 across all samples were retained. Differential expression analysis was performed on the raw counts of these retained genes using DESeq2 with a multifactorial design, which takes individual, organoid type, adjuvant type and interaction effect into account.

Raw sequencing reads in FASTQ format were assessed for quality using FastQC (v0.12.1). Adapter sequences and low-quality bases were removed with cutadapt[4] (v4.4). Filtered reads were aligned to the human reference genome GRCh38.p14 using STAR (v2.7.10b) with the corresponding NCBI RefSeq gene annotation file (GCF_000001405.40_GRCh38.p14_genomic.gtf) supplied during genome index generation and alignment. Gene-level read counts were obtained using featureCounts (v2.0.6) based on RefSeq annotation. Only uniquely mapped reads were retained for downstream quantification. The resulting raw count matrix was used for all downstream differential expression analyses.

Lowly expressed genes were filtered after transformation of raw counts using the rlog function in DESeq2. Genes with a 90th percentile of rlog-transformed expression greater than 1 across all samples were retained for downstream analyses. Differential expression analysis was performed on raw counts of retained genes using DESeq2 with a multifactor design including individual, organoid type, adjuvant, and their interaction. Unless otherwise specified, adjusted p values were controlled using the Benjamini–Hochberg procedure.

#### *Pathway Enrichment Analysis*

Gene Set Variation Analysis (GSVA) was conducted on the filtered rlog-transformed expression matrix using the GSVA R package. Reference gene sets included the MSigDB Hallmark and Reactome collections, immune-related Gene Ontology Biological Processes (GOBP), and Blood Transcriptomic Modules (BTMs) derived from human vaccine studies. To identify differentially enriched pathways, the resulting GSVA scores were analyzed using the limma package. A cell-means model was constructed by combining the antigen and adjuvant factors. To appropriately account for the repeated-measures design, intra-subject correlations were estimated using the duplicateCorrelation function (blocking by individual) and incorporated into the linear model, followed by empirical Bayes moderation (eBayes) to calculate statistical significance.

Over-representation analysis of differentially expressed genes was performed using the clusterProfiler[8] R package. Gene Ontology (GO) and Kyoto Encyclopedia of Genes and Genomes (KEGG) enrichment analyses were conducted on sets of significantly upregulated or downregulated genes, as specified for each comparison. Unless otherwise indicated, enrichment significance was evaluated using q values, with cutoff $q < 0.05$.

Gene set variation analysis (GSVA) was performed on the filtered rlog-transformed expression matrix using the GSVA R package[9]. Reference gene sets included MSigDB Hallmark, Reactome, immune-related Gene Ontology Biological Process terms, and Blood Transcriptional Modules (BTMs) derived from human vaccine studies[10]. Differential pathway enrichment was assessed from GSVA scores using limma[11]. A cell-means model was constructed by combining antigen and adjuvant conditions. To account for repeated measures, within-individual correlation was estimated using duplicateCorrelation with individual specified as the blocking factor and incorporated into the linear modeling framework. Statistical significance was calculated after empirical Bayes moderation using eBayes.

##### *Cell type deconvolution*

To estimate the relative abundance of distinct immune and stromal cell populations within our bulk RNA-seq cohort, we performed in silico cell type deconvolution using CIBERSORTx[12]. To ensure a highly accurate and biologically relevant deconvolution, we leveraged scRNA-seq dataset from the Human Tonsil Atlasas our reference transcriptome. Briefly, the annotated scRNA-seq count matrix, with the original cell type annotations, was obtained and utilized to construct a custom gene expression signature matrix according to CIBERSORTx's tutorials, as well as the bulk RNA seq matrix.

Cell type deconvolution of bulk RNA-seq data was performed using CIBERSORTx. We used an annotated single-cell RNA-seq dataset from the Human Tonsil Atlas[13] to generate a custom signature matrix following the CIBERSORTx workflow. Bulk RNA-seq expression profiles were deconvolved with the original cell type annotations of Human Tonsil Atlas according to CIBERSORTx's tutorials.

##### *Identification of conserved adjuvant-specific gene*

To identify candidate pan-antigen conserved adjuvant-response genes, we first fitted an additive model in DESeq2 (~ individual + antigen + adjuvant) to detect genes with a significant overall adjuvant effect after controlling for individual and antigen effects. Antigen-specific heterogeneity was then assessed by likelihood ratio testing of an interaction model (~ individual + antigen + adjuvant + antigen:adjuvant) against the additive model. Genes showing significant antigen-by-adjuvant interaction were excluded. Candidate conserved response genes were further required to show concordant effect directions across antigen strata.

##### *Quantification of GC-like zonation in organoid sections*

To quantify germinal center-like (GC-like) organization in organoid sections, CXCR4^high regions were operationally defined as dark zone-like (DZ-like) regions and CD83^high regions as light zone-like (LZ-like) regions, consistent with the analysis framework used in Fig. 1. For all analyses within a given experiment, images were processed using identical background-subtraction and thresholding settings in Fiji/ImageJ. GC-like regions of interest (ROIs) were manually defined on the basis of compact cellular aggregates containing spatially resolved CXCR4 and/or CD83 signal. Three complementary parameters were used to characterize GC-like zonation.

First, to assess functional polarization, the mean fluorescence intensity (MFI) of CXCR4 and CD83 was measured within each GC-like ROI, and a CXCR4/CD83 intensity ratio was calculated. This ratio was used to estimate the relative dominance of DZ-like versus LZ-like features within the same structure.

Second, to assess regional restructuring, area-fraction analysis was performed for each marker channel within the same GC-like ROI. Images were segmented using a standardized threshold applied identically across compared groups within the same experiment. The positive area for CXCR4 and CD83 was measured independently and expressed as a percentage of the total GC-like ROI area. Because CXCR4^high and CD83^high regions were quantified independently on the basis of marker-specific thresholded areas, DZ-like and LZ-like area fractions were not expected to sum to 100%. These measurements were used to compare the relative expansion or contraction of DZ-like and LZ-like compartments under different stimulation conditions.

For GC-like detectability analysis, each imaged field/well at each time point was scored as GC-like positive or negative. A GC-like-positive field/well was defined by the presence of a localized GC-like region showing discernible CXCR4-enriched and/or CD83-enriched organization. GC-like detectability was calculated as the number of GC-like-positive fields/wells divided by the total number of analyzed fields/wells at that time point and expressed as a percentage.

For representative spatial-segregation analysis, one GC-like ROI was selected from each displayed image when a detectable GC-like structure was present. The ROI was selected based on clear morphology and the presence of identifiable CXCR4-enriched and CD83-enriched domains. The Feret diameter of the GC-like ROI was measured in Fiji. A line of fixed width was then drawn across the major axis connecting the CXCR4-enriched and CD83-enriched regions, and fluorescence intensity profiles were extracted from the raw CXCR4 and CD83 single-channel images using the same line ROI. The dominant CXCR4 and CD83 peaks within the body of the GC-like ROI were identified from the line-scan profiles, excluding edge-associated fluctuations. Peak separation distance was calculated as the absolute distance between the dominant CXCR4 and CD83 peaks. Normalized CXCR4–CD83 peak separation was calculated by dividing the peak separation distance by the Feret diameter of the corresponding GC-like ROI. Images without detectable GC-like structures were recorded as not applicable for peak-separation analysis.

#### *Statistical analysis*

Statistical analyses were conducted in GraphPad Prism and/or R [4.2.3]. Data are presented as mean ± SEM. For comparisons between two groups, paired or unpaired two-tailed Student's t-tests were used as appropriate. For experiments involving two independent factors, such as time and treatment condition in time-course experiments with both unstimulated and stimulated groups, two-way repeated-measures ANOVA was used, followed by Tukey's multiple comparisons test for post hoc pairwise comparisons. For comparisons among multiple parallel groups at a single time point or under a single experimental condition, repeated-measures one-way ANOVA was used, followed by Dunnett's multiple comparisons test when each treatment group was compared with a shared control group. Statistical analyses were performed using R software. For comparisons between two groups, statistical significance was determined using the Wilcoxon test, with P-values adjusted for multiple testing via the Benjamini-Hochberg (BH) method to control the false discovery rate (FDR). To compare multiple treatment groups against a single control group while accounting for inter-subject variability, a Linear Mixed-Effects Model (LMM) was constructed using the lme4 package. In this model, the adjuvant type was treated as a fixed effect, and the individual subject was modeled as a random intercept to control for baseline variations. Subsequently, estimated marginal means (EMMs) were computed using the emmeans package to perform post-hoc pairwise comparisons between each adjuvant group and the blank control. P-values for these multiple comparisons were adjusted using Dunnett's method.

#### *Reference*


1. Wagar LE, Salahudeen A, Constantz CM, Wendel BS, Lyons MM, Mallajosyula V, et al. **Modeling human adaptive immune responses with tonsil organoids**. *Nature Medicine* 2021; 27(1):125-135.
2. Kim HK, Kim H, Lee MK, Choi WH, Jang Y, Shin JS, et al. **Generation of human tonsil epithelial organoids as an ex vivo model for SARS-CoV-2 infection**. *Biomaterials* 2022; 283:121460.
3. VanDussen KL, Sonnek NM, Stappenbeck TS. **L-WRN conditioned medium for gastrointestinal epithelial stem cell culture shows replicable batch-to-batch activity levels across multiple research teams**. *Stem Cell Research* 2019; 37:101430.
4. Martin M. **CUTADAPT removes adapter sequences from high-throughput sequencing reads**. *EMBnetjournal* 2011; 17.
5. Dobin A, Davis CA, Schlesinger F, Drenkow J, Zaleski C, Jha S, et al. **STAR: ultrafast universal RNA-seq aligner**. *Bioinformatics* 2013; 29(1):15-21.
6. Liao Y, Smyth GK, Shi W. **featureCounts: an efficient general purpose program for assigning sequence reads to genomic features**. *Bioinformatics* 2014; 30(7):923-930.
7. Love MI, Huber W, Anders S. **Moderated estimation of fold change and dispersion for RNA-seq data with DESeq2**. *Genome Biol* 2014; 15(12):550.
8. Yu G, Wang LG, Han Y, He QY. **clusterProfiler: an R package for comparing biological themes among gene clusters**. *OMICS* 2012; 16(5):284-287.
9. Hänzelmann S, Castelo R Fau - Guinney J, Guinney J. **GSVA: gene set variation analysis for microarray and RNA-seq data**. (1471-2105 (Electronic)).
10. Li S, Rouphael N, Duraisingham S, Romero-Steiner S, Presnell S, Davis C, et al. **Molecular signatures of antibody responses derived from a systems biology study of five human vaccines**. *Nat Immunol* 2014; 15(2):195-204.
11. Ritchie ME, Phipson B, Wu D, Hu Y, Law CW, Shi W, et al. **limma powers differential expression analyses for RNA-sequencing and microarray studies**. *Nucleic Acids Res* 2015; 43(7):e47.
12. Newman AM, Steen CB, Liu CL, Gentles AJ, Chaudhuri AA, Scherer F, et al. **Determining cell type abundance and expression from bulk tissues with digital cytometry**. *Nature Biotechnology* 2019; 37(7):773-782.
13. Massoni-Badosa R, Aguilar-Fernandez S, Nieto JC, Soler-Vila P, Elosua-Bayes M, Marchese D, et al. **An atlas of cells in the human tonsil**. *Immunity* 2024; 57(2):379-399 e318.
